# Force-mediated recruitment and reprogramming of healthy endothelial cells drive vascular lesion growth

**DOI:** 10.1101/2023.11.27.568780

**Authors:** Apeksha Shapeti, Jorge Barrasa-Fano, Abdel Rahman Abdel Fattah, Janne de Jong, José Antonio Sanz-Herrera, Mylène Pezet, Said Assou, Emilie de Vet, Seyed Ali Elahi, Adrian Ranga, Eva Faurobert, Hans Van Oosterwyck

## Abstract

Force-driven cellular interactions are known to play a critical role in cancer cell invasion, but have remained largely unexplored in the context of vascular abnormalities, partly due to a lack of suitable genetic and cellular models. One such vascular abnormality, cerebral cavernous malformation (CCM) is characterized by leaky, tumor-like vessels in the brain, where CCM mutant cells recruit wild-type cells from the surrounding endothelium to form mosaic lesions and promote lesion growth; however the mechanisms underlying this recruitment remain poorly understood. Here, we use 3D traction force microscopy in a in-vitro model of early angiogenic invasion to reveal that hyper-angiogenic CCM2-silenced endothelial cells enhance angiogenic invasion of neighboring wild-type cells through force and extracellular matrix-guided mechanisms. We show that mechanically hyperactive CCM2-silenced tips guide wild-type cells by exerting and transmitting pulling forces and by leaving degraded paths in the matrix as cues promoting invasion in a ROCKs-dependent manner. This transmission of forces is associated with a reinforcement of β1 integrin-dependent adhesive sites and actin cytoskeleton in the wild-type followers. We also show that during this process wild-type cells are reprogrammed into stalk cells through activation of matrisome and DNA replication programs, eventually leading to cell proliferation. These observations unveil a novel vascular lesion growth mechanism where CCM2 mutants hijack the function of wild-type cells to fuel CCM lesion growth. By integrating biophysical computational methodologies to quantify cellular forces with advanced molecular techniques, we provide new insights in the etiology of vascular malformations, and open up avenues to study the role of cell mechanics in tissue heterogeneity and disease progression.

## Introduction

Angiogenesis is emerging as a complex process regulated by not just biochemical signaling but also mechanical cues, with particular relevance for pathologies such as cancer^1–4^. Yet, how cellular forces drive aberrant angiogenesis in vascular diseases remains insufficiently explored. Cerebral Cavernous Malformations (CCM) is one such disease of the brain associated with regions of active angiogenesis^5^, and is caused by mutations in one of three CCM genes (CCM1/CCM2/CCM3)^6,7^. CCM lesions are thought to initiate through a cancer-like clonal proliferation of mutant endothelial cells (ECs)^8,9^. In advanced stages, symptomatic CCM lesions manifest as hemorrhagic, multicavernous structures comprising both mutant and wild-type (WT) ECs^10–14^. As these lesions expand, mutant cells attract WT ECs into the growing lesions^8,9^, making this step in CCM formation a crucial therapeutic target for reducing symptoms. The precise mechanisms underlying this recruitment remain unknown. Observations of dysregulated Rho-associated kinase (ROCK) signaling in CCM mutants, as well as the successful inhibition of symptomatic late stage lesions using Fasudil, a ROCK inhibitor, suggest a role for ROCK-dependent altered cell mechanics in lesion growth ^15–19^. However, nothing is known about the mechanical coupling between mutant and wild type cells, their surrounding extracellular matrix (ECM), and the consequences of these interactions on the fate of WT ECs during lesion growth.

Expanding upon the cancer-like mechanisms identified for CCM formation^20,21^, we asked if cooperative migration strategies such as those used by fibroblasts to regulate tumor cell invasion^22,23^ might also contribute to recruitment and reprogramming of WT ECs by CCM mutants. Since this multifactorial question requires an approach to systematically explore the mechanics of both cell-matrix and cell-cell interactions, we leveraged our three-dimensional (3D) biomimetic model of angiogenesis^24^ integrated with biophysical computations using 3D traction force microscopy and molecular investigations using single-cell RNA sequencing. Doing so, we discovered that hyper-angiogenic CCM2-silenced endothelial cells enhance angiogenic invasion of neighboring wild-type cells through force- and extracellular matrix-guided mechanisms. Mechanically hyperactive CCM2-silenced tips steer wild-type cells by exerting and transmitting pulling forces and leaving degraded paths in the matrix as cues to promote invasion in a ROCKs-dependent manner. Further, this recruitment of WT ECs involves their reprogramming into proliferative stalk cells characterized by upregulated expression of matrisome and DNA replication genes.

### Loss of CCM2 activates ROCK1-dependent filopodia formation in angiogenic tip cells

Human umbilical vein EC (HUVEC) monolayers were allowed to sprout into degradable, 3D polyethylene glycol (PEG) hydrogel based ECMs with sphingosine-1-phosphate (S1P) as a proangiogenic cue (Fig. 1a). siRNA silencing of CCM2 in EC monolayers previously showed increased angiogenic invasion in this model24. Given that filopodia are known to guide endothelial tip cells towards angiogenic stimuli^25,26^, we wondered if these structures were altered in response to CCM2 loss. Upon closer examination, we observed that highly angiogenic CCM2-silenced sprouts, referred to as siCCM2 ECs, (Fig. 1b), exhibited denser localization of F-actin (Fig. 1b,c) and a pronounced branchy tip cell phenotype^26^ characterized by longer filopodia compared to control sprouts, denoted as siCT (Fig. 1b,d). Additionally, siCCM2 ECs displayed hypersensitivity to angiogenic signaling, demonstrated by their increased sprouting even at lower S1P concentrations, while siCT ECs rarely sprouted under these conditions (Extended Data Fig. 1-A). Notably, siCCM2 ECs also exhibited an enhanced ability to sense the matrix as evidenced by their extension of numerous small filopodia into a non-degradable PEG-based matrix, a behavior not observed in siCT ECs (Extended Data Fig. 1-B). Given that Rho/ROCKs (Rho-associated protein kinase)-dependent cytoskeletal remodeling differentially regulates invasion upon CCM2 loss^24,27^, we further explored the sprouting behavior of siCCM2 ECs following additional silencing of ROCK1 (siCCM2+siROCK1) or ROCK2 (siCCM2+siROCK2). The hyperactivated filopodial matrix-sensing phenotype of CCM2 mutant ECs was rescued by the additional silencing of ROCK1 but not ROCK2 (Fig.1b,c), consistent with the rescue of invasiveness observed only with ROCK1 (Fig. 1b). These findings demonstrate that only ROCK1 is necessary for filopodia formation, reminiscent of its role in the generation of dendritic spines through the segregation of activated Rac1^28^.

**Figure 1.**
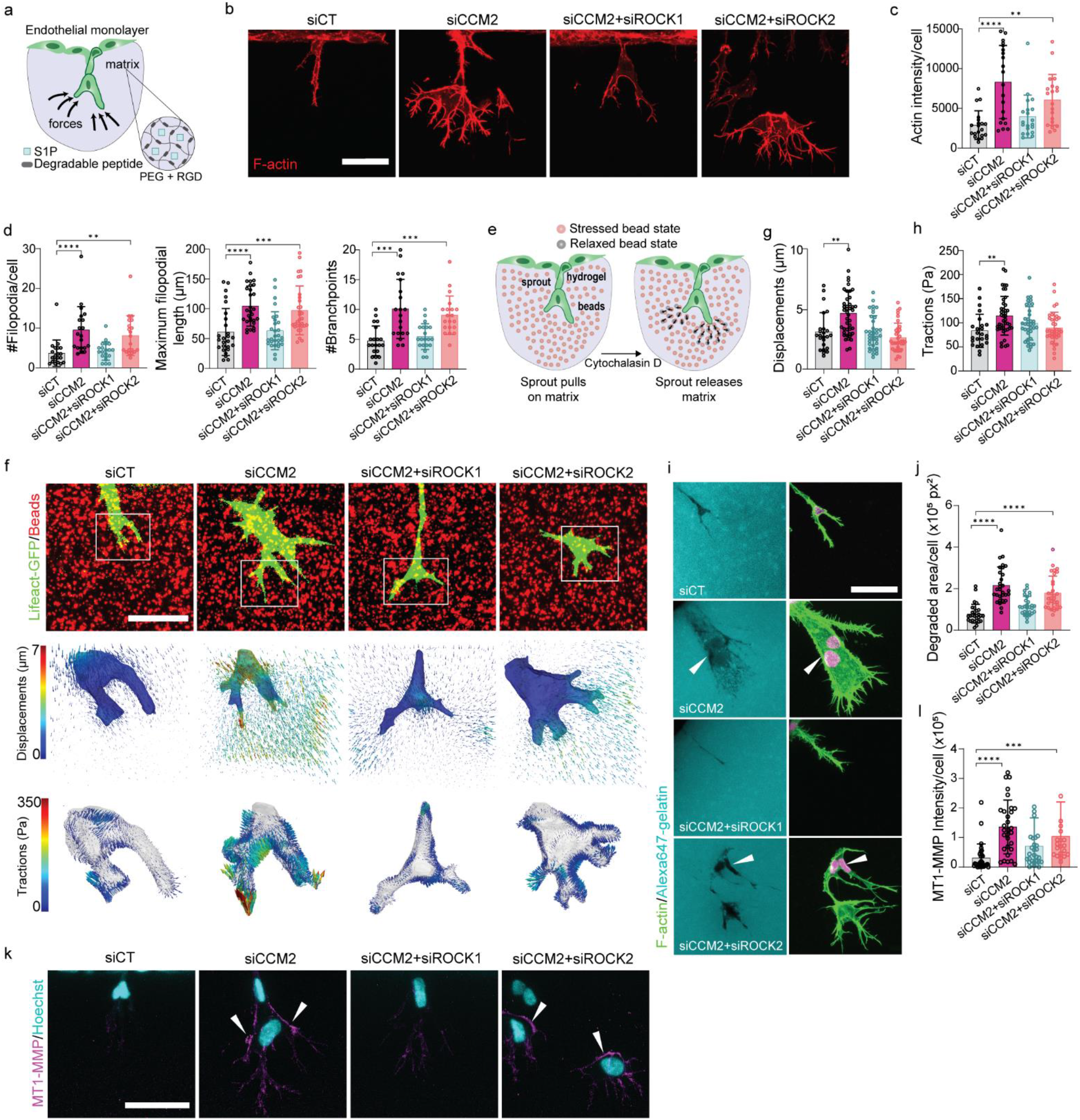
Enhanced angiogenic sprouting in CCM2-silenced endothelial cells relies on ROCKs-dependent force generation and ECM degradation. **a.** Schematic representation of the PEG hydrogel based 3D angiogenic invasion assay. **b.** Maximum intensity projections of F-actin expression after overnight sprouting of ECs transfected with siRNAs. **c.** Quantification of the total actin intensity in leading tip regions (n=20, for all conditions). **d.** Filopodial morphological metrics in leading tip regions (number of filopodia (2 independent trials; n=20 for all conditions), maximum filopodial length (2 independent trials; siCT, n=26; siCCM2, siCCM2+ROCK1, n=29; siCCM2+ROCK2, n=31), number of branchpoints in each tip (2 independent trials; n=20 for all conditions)). **e.** Schematic representation of 3D traction force microscopy on angiogenic sprouts. **f.** Representative maximally projected images of beads (in red) overlaid with GFP-labeled sprouts (top) and the corresponding 3D renders of displacements (middle) and traction fields (bottom) computed around angiogenic sprout tips of silenced ECs after 18h of invasion. **g,h.** Automated quantification of the 90^th^ percentile bead displacements (**g,** in µm) and tractions (**h,** in Pa) exerted by force exerting sprout tip regions (3 independent trials; siCT, n=24; siCCM2, n=43; siCCM2+ROCK1, n=37; siCCM2+ROCK2, n=35). **i.** Representative minimum intensity projections of fluorescent gelatin degraded by silenced ECs during 18h of invasion (left) and the corresponding sprouts labeled with f-actin (green) and nuclei with Hoechst (magenta) (right). White arrows showing higher degradation under the cell body in the rear. **j.** Automated quantification of total area of gelatin degradation normalized to the number of nuclei (pixel^2^; 3 independent trials; siCT, n=28; siCCM2, n=30; siCCM2+ROCK1, n=29; siCCM2+ROCK2, n=30) **k.** Representative maximally projected immunofluorescence images of MT1-MMP expression (magenta) and nuclei stained with Hoechst (cyan). **l.** Automated quantification of total MT1-MMP intensity per nuclei computed in maximum intensity projections (3 independent trials; siCT, siCCM2, siCCM2+ROCK1, n=30; siCCM2+ROCK2, n=20, presented here as averages after normalizing to the control of each trial). All scale bars are 50µm. Error bars represent SEM Kruskal-Wallis significance test with Dunn’s correction for multiple comparisons with the control, p-value <0.05(*) <0.01(**) <0.001(***).

### Loss of CCM2 enhances cell-ECM forces and matrix degradation through ROCKs

Collective angiogenic invasion requires complex sprout-ECM interactions and active cellular machinery for applying mechanical force^29,30^. Given that inhibition of actomyosin contractility with blebbistatin treatment reduces sprouting of siCCM2 ECs^24^, we hypothesized that the enhanced invasion accompanied by a hyperactivated tip phenotype of CCM2 mutant ECs must also translate to higher forces on the matrix. To test this, we incorporated fluorescent beads into soft (0.5kPa, Extended Data Fig. 2-A), PEG-based ECMs as markers for measuring cell-induced hydrogel deformations during angiogenic sprouting using 3D traction force microscopy (TFM, Fig. 1e). As predicted, siCCM2 sprout tips pulled more strongly on the ECM to generate 35-40% higher bead displacements and subsequent tractions on the ECM compared to siCT sprout tips (Fig. 1f-h). Additional silencing of ROCK1 reverted the increase in ECM displacements and tractions, consistent with the decrease in filopodia upon ROCK1 depletion (Fig. 1d,f). Taken together, this suggests that tractions exerted by siCCM2 ECs were associated with the abundant ROCK1-dependent tip cell filopodia. Surprisingly, the numerous filopodia still observed in siCCM2+siROCK2 sprouts did not translate to higher forces on the ECM (Fig 1d,e). This suggests that ROCK2 is necessary for force generation by actin-rich filopodia, as also reported for contractile forces generated by dendritic spines^28^. However, a complete restoration of the increased forces observed upon CCM2 loss seemed contradictory with the sustained increase in invasiveness and F-actin of these siCCM2+siROCK2 sprouts (Fig 1b).

We wondered if matrix degradation was an additional factor contributing to this increase in invasion. To answer this, we combined PEG matrices with fluorescently labeled gelatin to visualize matrix degradation. Loss of CCM2 increased ECM degradation in 3D during angiogenic invasion (Fig 1h, i), consistent with previous observations on 2D matrices^24^. ECM degradation was rarely noted in filopodial tip regions and was higher at the rear of the cell in the vicinity of the nuclei (Fig. 1i, white arrows), as is also reported during confined tumor migration^31^. Immunostaining for MT1-MMP also confirmed an increase in MMP expression, especially at the rear of the cell (Fig. 1k,l). siCCM2 sprouts additionally silenced for ROCK2 also showed higher matrix degradation which correlated with their increase in invasion (Fig. 1i,j). Only ROCK1 but not ROCK2 silencing restored MT1-MMP expression and ECM degradation. Subsequently, treating siCCM2 ECs with GM6001 (broad spectrum MMP inhibitor, MMPi) also reduced invasion (Extended Data Fig. 2B). Timelapse imaging of the matrix during siCCM2 sprouting showed that degradation did not co-localize with the force exerting tip regions, consistent with previously shown matrix degradation at sprout stalks^32^, thereby confirming it was not the driver for higher bead displacements (Supplementary Video 1).

Combined, these data attribute the enhanced angiogenic invasion upon CCM2 loss to a combination of hyperactivated filopodia of CCM2 mutant tip cells at the front, increased pulling forces by highly matrix-sensing, and MMP-driven matrix degradation at the rear aiding migration of the cell body. We noted that an increase in cell-ECM forces upon CCM2 loss requires both ROCK1 and ROCK2, while degradation is solely mediated by ROCK1. Above all, we confirmed that dysregulated mechanics is crucial to the overactivation of sprouting angiogenesis upon CCM2 loss.

### Hyper-angiogenic CCM2-silenced EC tips lead and stimulate wild-type EC invasion

We then turned our attention to the concept of mosaicism in CCM lesions evidenced by the presence of not just mutant, but also wild-type cells likely recruited from the healthy endothelium^8,9^. We have previously shown that siCCM2 ECs use a senescence-associated secretory phenotype (SASP) to attract surrounding wild-type (WT) ECs^24^. Moreover, cancer cells undergoing SASP aid tumor development by promoting tumor angiogenesis and tumor cell invasion^33^. Theorizing that CCM progression is driven by CCM2 mutants similarly activating angiogenic invasion in WT ECs, we questioned the underlying mechanism of such recruitment. To first assess the role of putative paracrine factors secreted by siCCM2 ECs on WT ECs, we combined PEG matrices with media conditioned by siCCM2 or siCT ECs for 24h. We observed no significant paracrine angiogenic effects on invasion (Extended Data Fig. 3-A), suggesting that the recruitment of WT ECs observed in our in-vitro model was likely due to physical mechanisms of cooperative invasion.

To further explore angiogenic activation of WT ECs by CCM mutants, we visualized collective sprouting of LifeAct-RFP tagged WT ECs and LifeAct-GFP tagged, silenced ECs (CT or CCM2) mixed to form mosaic sprouts at varying percentages of silenced ECs (10%, 20% or 50%) (Fig. 2a). We noted an increase in invasion of WT ECs in CCM2 mutant mosaics composed of 50% and 20% siCCM2 ECs, but not at 10% (Fig. 2b). This implied that a minimum critical mass of siCCM2 ECs was necessary in our model to modify angiogenic behavior of neighboring WT ECs. This is in line with in-vivo models of CCM pathogenesis and human lesions where only between 15 to 20% of ECs carry an actual CCM mutation^8,12–14^.

**Figure 2.**
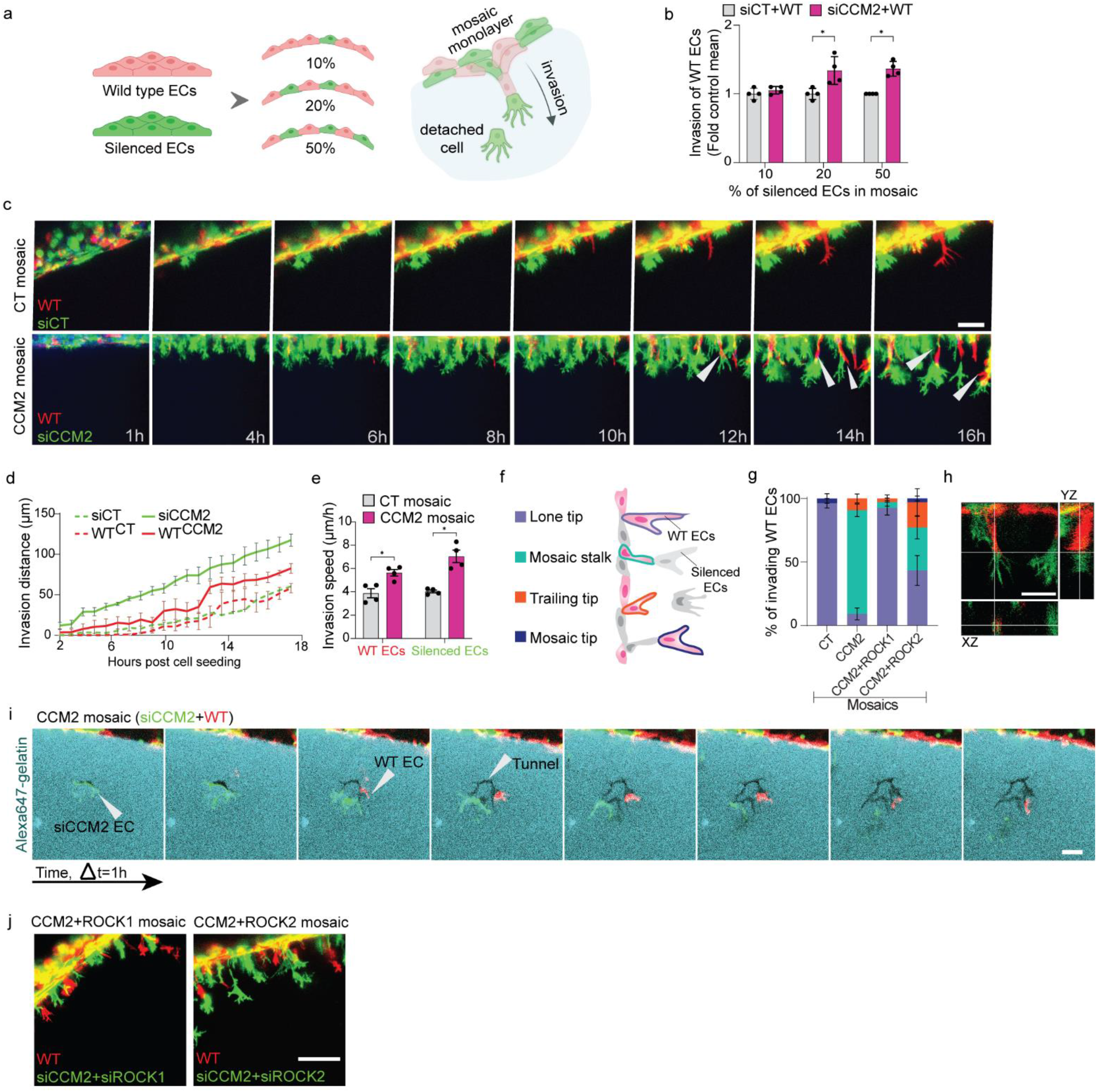
Hyper-angiogenic siCCM2 ECs stimulate invasion of wild-type ECs by leading mosaic sprouting in a ROCKs-dependent manner. **a.** Schematic representation of mosaic invasion assay combining Lifeact-RFP labeled wild-type (red) with Lifeact-GFP labeled (green) ECs siRNA silenced for Control, CCM2, CCM2+ROCK1 or CCM2+ROCK2. **b.** Maximum invasion distance of WT ECs in CT mosaics (siCT+WT) versus CCM2 mutant mosaics (siCCM2+WT) at varying percentages of silenced ECs (n=4, 35-40 positions/condition, quantified as averages after normalizing to the control for each trial). **c.** Maximally projected confocal images captured during 18h-invasion of 1:1 mixtures of WT ECs (in red) and siCT or siCCM2 ECs (in green). siCCM2 ECs lead WTs during invasion (white arrows). These images are representative of 12 FOVs from 4 independent trials (scale bar, 50 µm). **d.** Automated quantification of average invasion distance of WT and silenced ECs in CT and CCM2 mutant mosaics over time (n=3, 9 FOVs/condition). **e.** Quantification of invasion speed (µm/h) of WT or silenced ECs in CT and CCM2 mutant mosaics (n=4, averages based on 12 positions/condition) **f.** Schematic representation of categories of wild-type ECs defined based on their position relative to silenced ECs at the start of ECM invasion. **g.** Quantification of the percent categorical distribution of the various positional modes of invasion described (in **g**) for WT ECs in CT versus CCM2 mutant mosaics. **h.** An XZ and YZ section of CCM2 mutant mosaic sprout showing contact between WT (in red) and siCCM2 (in green) ECs (scale bar is 50 µm). **i.** Confocal images taken at 1h intervals during overnight invasion of a 1:1 mix of WT and siCCM2 ECs. Composite images are composed of a single z-plane of fluorescently labeled gelatin combined with the two cell types. Follower WT EC modifies direction of migration by sensing tunnels created by leading siCCM2 EC (scale bar, 50 µm). **j.** Maximally projected confocal images after 16h-invasion of 1:1 mixtures of WT ECs and siCCM2 ECs additionally silenced for either ROCK1 or ROCK2 (scale bar, 100µm). Error bars represent SEM Kruskal-Wallis significance test with Dunn’s correction for multiple comparisons with the control, p-value <0.05(*) <0.01(**) <0.001(***).

To focus our efforts on understanding WT EC recruitment, we used a 1:1 mix of WT and silenced ECs for all following mosaic studies. Tracking dynamics of angiogenic sprouting over 18h showed that both siCCM2 and WT ECs begin invading earlier and faster in CCM2 mutant mosaics than in CT mosaics (Fig. 2c-e, Supplementary Videos 2A-B). However, siCCM2 ECs initiated invasion earlier than the WT ECs in these CCM2 mutant mosaics and had an invasive advantage as shown by their increased invasion speed (Fig. 2d,e). We identified four invasion modes of WT ECs based on their positions in relation to siCCM2 or siCT ECs at the start of ECM invasion (Fig. 2f). WTs began invading the matrix as: leading tips (“lone tips”), leading tips in contact with a following silenced stalk EC (“mosaic tips”), following stalks in contact with a leading silenced EC (“mosaic stalks”), or tip cell trailing behind a distant silenced EC (“trailing tips”). Remarkably, we observed that 80% of invading WT ECs in CCM2 mutant mosaics begin doing so in “mosaic stalk” positions in direct contact with a leading siCCM2 EC in front at the tip of the angiogenic sprout (Fig. 2c,g,h). In contrast, WT ECs in CT mosaics begin invading the matrix mainly as “lone tips” and never assume “mosaic stalk” positions (Fig. 2c,g). We concluded that CCM2 mutant ECs are capable of inhibiting a tip phenotype in WT ECs.

### ROCKs dependent forces and ROCK1-dependent tunnels guide mosaic intercellular communication

A small percentage of the sprouting WT ECs in CCM2 mutant mosaics (10%, Fig. 2f,g) began invading the matrix without any direct contact based communication as “trailing tips” behind distant leading siCCM2 ECs. Dynamic co-visualization of matrix degradation and migration occasionally showed trailing WT ECs modify their direction of migration to turn towards tunnels created by distant leading mutant ECs (Fig. 2i, Supplementary Video 1). This combined with the increase in matrix degradation during mosaic sprouting of siCCM2 and WT ECs (Extended Data Fig. 4-A, B) indicated that, in a subset of cases, reduced physical impedance and the likely residual, fragmented ECM components in empty tunnels could serve as a recruitment cue for contact-free guidance of invading WT ECs.

Strikingly, WT ECs in mutant mosaics regain tip cell behavior when either ROCK1 or ROCK2 is depleted in the siCCM2 EC (Fig. 2g, j), suggesting that the invasive advantage of CCM2 mutants over WT ECs might result from their ability to exert higher tractions (Fig. 1h). Further, ROCK1 depletion was more efficient than ROCK2 in counteracting tip cell inhibition of WT ECs (Fig. 2g). This indicates that ROCK1-dependent matrix degradation contributes to the invasive advantage of CCM2 mutants (Extended Data Fig. 4-A, B). Since these hyper-degradative skills were restored by both blebbistatin and MMPi treatment, aberrant actomyosin contractility again proved necessary for increased degradation during mosaic sprouting in 3D (Extended Data Fig. 4-A, B). Blocking myosin contractility with blebbistatin or MT1-MMP-driven degradation with MMPi restored the increase in invasion of both WT and siCCM2 ECs in mosaics confirming that these mechanisms were necessary (Extended Data Fig. 4-C, D). To exclude biochemical signaling through Notch to be a factor in the enhanced invasion of WTs, we evaluated DAPT (indirect inhibitor of Notch) treated CCM2 mutant mosaic sprouting and noted no change in invasion of WT ECs (Extended Data Fig. 4-C, D). Taken together, we conclude that siCCM2 ECs lead angiogenic invasion by imposing a ROCK-dependent, contact-guided stalk position in migrating WT ECs. For the first time, this provides evidence for a role of cell mechanics in CCM lesion mosaicism.

### Mechanically hyperactive CCM2-silenced tips steer passive WT ECs through pulling forces and reinforce their actin cytoskeleton

Building upon the coherent increase in force exertion and invasion observed for siCCM2 EC-only sprouts in Fig. 1f, we asked if the increase in invasion of WT ECs in the presence of siCCM2 ECs (Fig. 2c,d) was also accompanied by higher forces on the matrix. To answer this, we used 3D TFM to quantify force-induced matrix displacements generated by both cell populations after overnight mosaic invasion (Fig. 3a,b). Analyzing solely cell-associated matrix displacements (Fig. 3c), we discovered that a significantly lower percentage of invading WT ECs actively exerted forces on the matrix in CCM2 mutant versus CT mosaics (Fig. 3d). This was in line with their stalk cells-like behavior (Fig. 2g), which are shown to exert only low contractile forces on the matrix^34^. Interestingly, depletion of either one of the ROCKs rescued the percentage of force exerting WTs (Fig. 3d), consistent with their rescue as tip cells (Fig. 2g, j). However, the magnitude of matrix displacements generated by the force exerting WT ECs did not vary between the four conditions of cell mosaics and remained similar to control levels (Fig. 3e). Meanwhile, the percentage of force-exerting silenced ECs in mosaic sprouts was drastically higher for the hyper angiogenic tip-like siCCM2 ECs than siCT ECs, and was not modulated by ROCKs (Extended Data Fig. 5-A). Their increase in force exertion was however ROCKs-dependent in agreement with our findings during non-mosaic invasion (Fig. 3f, Fig. 1f,h). Taken together, this establishes that CCM2 loss drives a ROCKs-dependent non-autonomous inhibition of leader cell behavior in WT ECs, preventing them from transmitting forces to the matrix. This inhibition depends on the increased capacity of siCCM2 tip cells to generate forces. Above all, cell-ECM forces of WT ECs did not increase and could not explain their increased invasiveness in CCM2 mutant mosaics.

**Figure 3.**
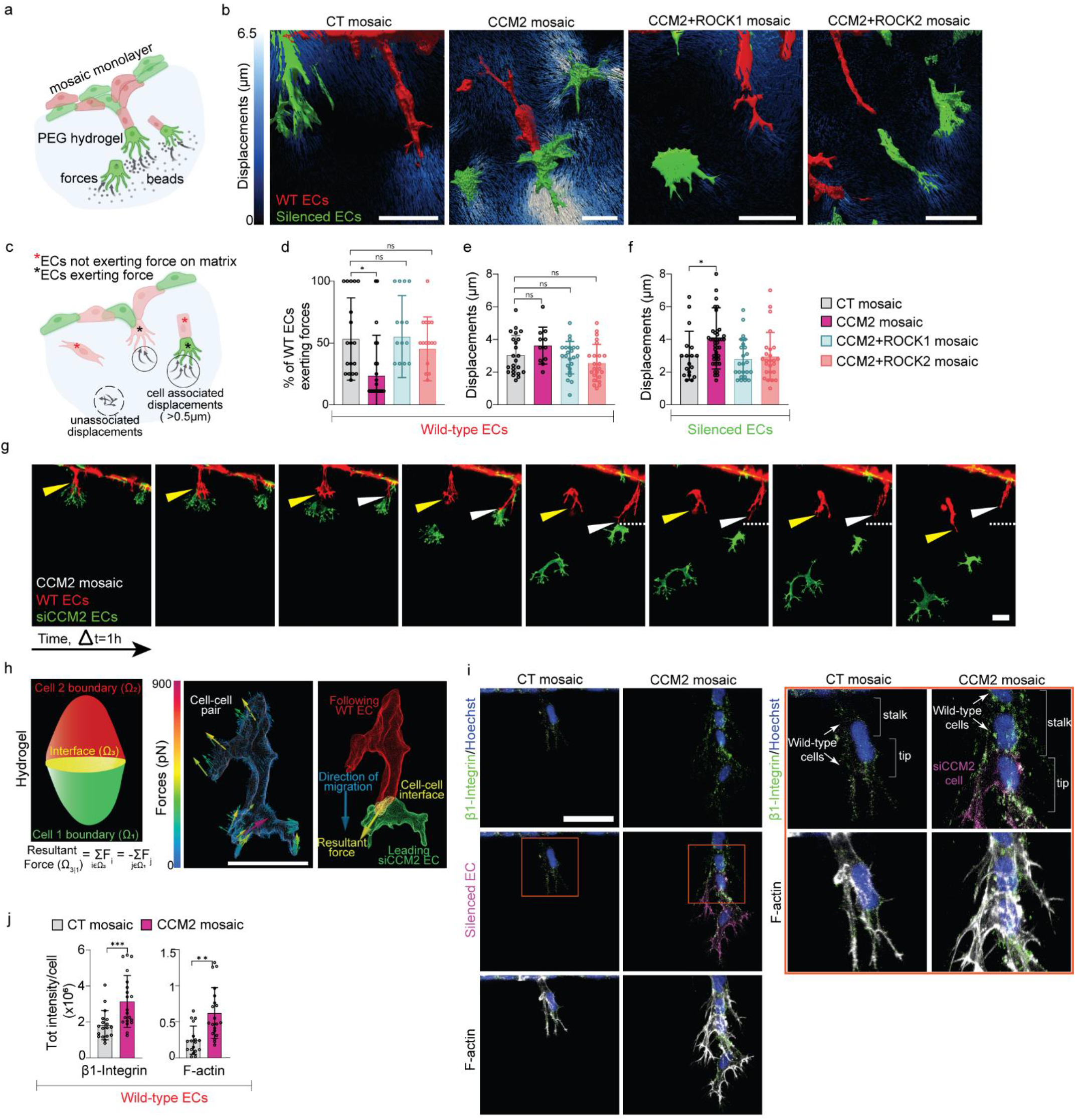
Mechanically hyperactive CCM2-silenced tips steer passive WT ECs through pulling forces and reinforce their β1 integrin adhesions and actin cytoskeleton. **a.** Schematic representation of 3D TFM compatible mosaic invasion assay embedded with fluorescent beads. All experiments used a 1:1 mixture of wild-type (red) and siRNA silenced ECs (green; CT, CCM2, CCM2+ROCK1 or CCM2+ROCK2). **b.** Representative 3D renders of displacements (in µm) computed around mosaic angiogenic sprouts imaged after overnight invasion. **c.** Schematic illustrating the criterion imposed for quantifying the fraction of ECs actively exerting pulling forces on the matrix. Only displacements larger than 0.5 µm exerted clearly localized in the vicinity of the cell were considered to be cell associated active pulling on the matrix. **d.** Quantification of the percentage of WT ECs actively exerting forces on the matrix based on criteria in (**c**). 3 independent trials; CT mosaic, n=19; CCM2, n=21; siCCM2+ROCK1, n=16; siCCM2+ROCK2, n=17. **e,f.** Maximum pulling displacements (µm) by force exerting WT ECs (**e,** 3 independent trials; CT mosaic, n=23; CCM2, n=11; siCCM2+ROCK1, siCCM2+ROCK2, n=23) or silenced ECs (**f,** 3 independent trials; CT mosaic, n=18; CCM2, n=44; siCCM2+ROCK1, n=24; siCCM2+ROCK2, n=25). **g.** Maximum intensity projection of confocal images taken at 50 min intervals during overnight invasion of a 1:1 mix of WT (red) and siCCM2 (green) ECs. Follower WT ECs decelerate invasion (yellow arrow) or retract (white arrow, dotted line showing migration front) upon losing contact with hyperactive leading siCCM2 ECs. **h.** (**left**) Schematic of a mosaic cell pair depicting the two cell-hydrogel interfaces (green, Ω_1_; red, Ω_2_), the cell-cell interface (yellow, Ω_3_), forces exerted by the cells on the hydrogel (**F**), and the force balance equation to quantify resultant force on the interface. (**middle**) Force exerted by cells on the ECM for a WT and siCCM2 mosaic cell pair. (**right**) The resultant non-zero force vector exerted by the leading cell on the cell-cell interface (yellow arrow) and the direction of migration (blue arrow). **i**. Representative confocal immunofluorescence images (maximum projections) of (**top**) β1 integrin (green) and Hoechst-labeled nuclei (blue), (**middle**) fluorescently labeled silenced EC (magenta) and (**bottom**) F-actin (gray). Cells not labeled in magenta are WT ECs. **i.** Zoom in of contact with stalk WT ECs. All scale bars are 50µm. **j.** Automated total intensity quantification of β1 integrin expression and F-actin measured in regions containing only wild type ECs cropped from maximum projection images and normalized to the number of nuclei (2 independent trials; CT mosaic, n=18; CCM2 mosaic, n=20). Error bars represent SEM; one-way ANOVA Kruskal-Wallis significance test with Dunn’s correction for multiple comparisons with the control, p-value <0.05(*) <0.01(**) <0.001(***).

Cancer-associated fibroblasts have been shown to guide cancer cell invasion through intercellular force transmission^22^. We wondered if a similar mechanical communication could explain the collective migration of non-force exerting WT stalks behind highly force exerting siCCM2 tips. When observing invasion dynamics, we often noted that hyper-migratory siCCM2 tip cells detached and invaded farther away from a WT stalk leaving it behind (Fig. 2c, 3g, Supplementary Video 3). Interestingly, the WT stalk seemed to either retract (Fig. 3g, white arrow) or decelerate migration (Fig. 3g, yellow arrow) after detachment of the siCCM2 tip cell. This suggested that WT ECs were likely pulled forward by the leading siCCM2 tip. Therefore, to evaluate if leading siCCM2 ECs were mechanically interacting with stalk WT ECs at cell-cell contacts, we developed a novel 3D TFM approach based on first principles, i.e. linear and angular momentum conservation^35^, to quantify the resultant (net) force at this interface (Fig. 3h, Supplementary Note 2). By recovering the forces exerted on the ECM by a detached cell pair of a siCCM2 leader and a WT follower, we found that the average cell-ECM traction exerted by the mutant tip largely exceeded that by the WT stalk (Fig. 3h, Table 1 in Supplementary Note 2). Further, quantifying the resultant cell-cell force vector at the interface between these cells showed a substantial, non-zero force (∼14 nN) and corresponding average cell-cell traction (∼7 pN/µm^2^), indicating that the two cell types mechanically interact by means of pulling forces. Given that the resultant force from the tip cell on the stalk cell was aligned with the direction of migration of the cell pair (see Fig 3h), we deduced that these cell-cell pulling forces are important for bringing the WT stalk forward and to ensure collective migration of the tip and stalk cell (see detailed interpretation presented in Supplementary Note 2). Taken together, we concluded that siCCM2 tip ECs are capable of mechanically steering the invasion of passive WT stalks through the combined action of tip cell-ECM pulling forces to direct the sprout tip, and tip cell-stalk cell pulling force transmission to direct the sprout stalk.

Remarkably, these pulling forces experienced by WT ECs were further associated with a reinforcement of the actin cytoskeleton, as evidenced by an increase in β1 integrin-dependent focal adhesions (FA) and actin stress fibers transduced along the follower WT ECs (Fig. 3i, j). Despite the lower percentage of mechanically active WT ECs, β1 integrin-dependent cell-matrix contacts were elevated in WT ECs from CCM2 mutant mosaics. Interestingly, this cytoskeleton reinforcement was only dependent on ROCK1 but not ROCK2, consistent with the specific role of ROCK1 in the formation of β1 integrin-anchored stress fibers at the rear of migrating cells (Extended Data Fig. 5-B, C)^36^. We concluded that the mechanically hyperactive siCCM2 tip cells stimulate actin cytoskeleton reinforcement of the passive WT stalk cells to encourage their invasion and speculated on a mechanism involving pulling forces at cell-cell junctions.

### Single-cell RNA sequencing reveals an in vivo-like angiogenic transcriptomic program in sprouting CCM2-silenced ECs

Contextual mechanical forces experienced by cells from mechanosensing interactions with the ECM, or through force transmission from adjacent cells, have a considerable influence on gene expression programs as demonstrated for stem cell fate, tissue morphogenesis and collective cell migration^37–40^. We, therefore, asked if the dysregulated mechanics of siCCM2 ECs, and the subsequent exposure of neighboring WT ECs to abnormal mechanical interactions, mediated transcriptomic changes in WT ECs. To investigate this, we performed single cell RNA sequencing (scRNA-seq) on FACS-sorted WT and siCCM2 ECs 24h after invasion in either CT or CCM2 mutant mosaic conditions (Fig. 4a). WT cells isolated from CT mosaics were annotated as WT, and those from CCM2 mutant mosaics as WT-CCM2. The dataset was equally distributed between WT (32%), WT-CCM2 (32%) and siCCM2 (36%) conditions (Fig. 4b). However, a clear segregation was observed between siCCM2 clusters and both the WT/WT-CCM2 clusters confirming the evident effect of CCM2 silencing on the transcriptomic profile of the mutant ECs (Fig. 4b). Graph-based clustering of the 1434 ECs retained for analysis identified seven clusters based on upregulated genes associated with biological functions related to proliferation, senescence, and angiogenesis (clusters angiogenesis1/2/3) as well as two additional clusters with genes related to vasculature remodeling and RNA splicing (Fig. 4c, Supplementary file 1). The WT and WT-CCM2 conditions largely comprised proliferative, blood vessel remodeling and angiogenesis regulating cells, whereas loss of CCM2 prompted migration and sprouting angiogenesis activity, as well as non-proliferative G1 arrest, consistent with our prior discovery of SASP in CCM^24,41^. To corroborate the biological relevance of the siCCM2 ECs derived from our 3D in-vitro model of angiogenic invasion, we sought to match their dysregulated genes with previously established transcriptomic signatures of CCMs in vivo. siCCM2 ECs indeed showed a clear upregulation of signaling pathways known to play a pivotal role in lesion progression, such as TGF-beta, p53, nuclear beta catenin, inflammation, endothelial to mesenchymal transition, upregulation by RhoA and angiogenesis (Fig. 4d, Supplementary file 2).

**Figure 4.**
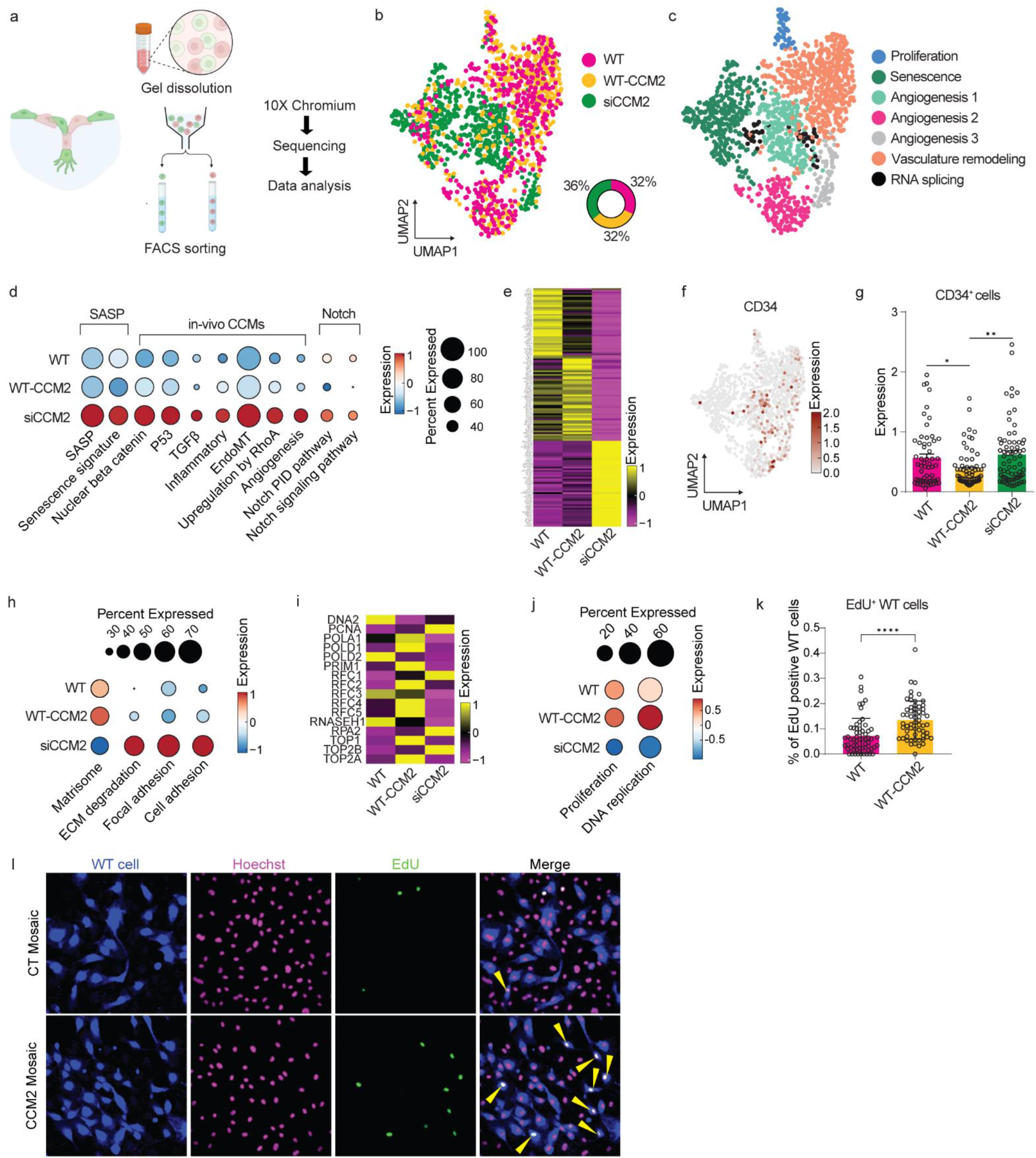
Single-cell RNA sequencing reveals a CCM2 EC mediated reprogramming of WT ECs into a stalk cell identity. **a.** Schematic representation of the single-cell RNA-sequencing workflow. **b.** UMAP and distribution of the combined dataset (WT, WT-CCM2 and siCCM2 ECs) **c.** UMAP of the combined dataset annotated with identified clusters of upregulated genes associated with proliferation, senescence, angiogenesis (angiogenesis 1, 2, 3), vasculature remodeling and RNA splicing (Supplementary file 1). **d.** Differential expression of major signaling pathways genes known to play a crucial role in CCMs in-vivo and in-vitro, shown in a dotplot. **e.** Heatmap of top 50 differentially expressed genes for each cell dataset. **f.** UMAP of the combined dataset showing CD34 positive tip cells. **g.** Quantification of normalized CD34 expression in CD34+ cells in different conditions. Unpaired two-sided t-test with exact p-values of 0.008 (**) and 0.0008 (***). **h.** Dotplot representation of matrisome, ECM degradation, focal adhesion and cell adhesion signaling pathways compared across the dataset. **i.** Heatmap showing the expression of genes associated with DNA replication and proliferation. j. Dotpot representation for comparison of DNA replication and proliferation gene expression between cell types. **k.** Quantification of the fraction of edU positive wild type ECs in 2D mosaics of WT and silenced (siCT or siCCM2) ECs 48h after seeding. Mann-Whitney test with p-value <0.0001 (3 independent trials; WT, n=60; WT-CCM2, n=59). **l.** Representative confocal images of fluorescently labeled wild-type ECs (blue), Hoechst-labeled nuclei (pink), EdU-positive nuclei (green) and the merged image. Nuclei in white upon merge represent EdU positive wild-type ECs (see yellow arrows).

### Recruitment of WT ECs by CCM2-silenced leaders induces reprogramming into a proliferative stalk cell identity

Beyond their phenotypic change observed during mosaic invasion with siCCM2 ECs (Fig.3h,i), we wondered if WT ECs cells undergo changes in their gene expression program. To address this, we further compared the transcriptomic landscape of WT versus WT-CCM2. Heatmaps of top 50 differentially expressed genes (DEGs) showed a distinct heterogeneity in the transcriptional signatures of the three conditions (Fig. 4e, Supplementary file 3). While siCCM2s showed the most pronounced differences as expected, WT-CCM2 signatures discernibly deviated from those of WT. Selecting for CD34+ cells^42^, we confirmed a significant increase in tip cells for siCCM2 ECs, alongside a distinct decrease in tip cells for WT-CCM2 relative to WT (Fig. 4f, g). Consistent with their tip cell identities characterized by high force exertion (Fig. 3b), siCCM2 ECs displayed elevated expression of genes associated with ECM degradation, focal adhesions, and cell adhesion (Fig. 4h).

In contrast, the predominantly stalk WT-CCM2s presented upregulated genes linked to ECM deposition termed matrisome (Fig. 4h, Supplementary file 4). Others have identified an upregulated matrisome signature underlying the angiogenic switch of normal cells during tumorigenesis, as well as matrix adhesion being an adaptive response driving cancer aggressiveness^43–45^. Further, increased mutant proliferation is typically associated with CCM growth^20^, a concept seemingly at odds with the non-proliferative, senescent nature of siCCM2 ECs in our in-vitro model. We wondered if, in conjunction with activated matrisome-driven signaling, WT-CCM2 may contribute to proliferation-driven CCM growth in line with their stalk identities. Strikingly, when compared with WT and siCCM2 ECs, WT-CCM showed a distinct upregulation of DNA replication and proliferation pathways (Fig. 4i,j). To validate this finding, we first checked for EdU-positive WT nuclei in CT or CCM2 mutant mosaics after 24h of invasion and observed no differences. We then explored this in 2D by comparing EdU-positive WT nuclei from a 1:1 mosaic with siCCM2 versus siCT ECs cultured for 24h or 48h. Notably, we observed a significant increase in EdU positive WT nuclei in CCM2 mutant compared to CT mosaics after 48h in 2D (Fig. 4k), but saw no clear differences at the 24h timepoint (Extended Data Fig. 6-A). This suggested that our 24h sprouting assay may have limitations in exploring this mechanism in the 3D context, despite the upregulation of DNA replication genes observed by single cell RNA sequencing. In summary, this demonstrated that mechanically dysregulated siCCM2 ECs are capable of upregulating stalk-like matrix deposition and proliferation programs in wild-type ECs.

## Discussion

Several recent discoveries have significantly advanced our understanding of CCM lesion pathogenesis through genetic and molecular investigations^7,10,20,46^. Nevertheless, several key questions regarding the role of wild-type endothelial cells in fueling CCM lesion growth and their non-autonomous activation by CCM mutants remain unanswered. The most noteworthy conceptual advance of our study is the establishment of a novel model in which CCM mutants use force- and ECM-mediated physical mechanisms to attract and reprogram wild-type cells from the healthy endothelium, potentially exacerbating lesion growth (Fig.5). This cooperative invasion strategy involving force transmission and microenvironmental cues bears striking resemblance to mechanisms observed in cancer progression^22,23,47^. The highly contractile, tip-like, leading CCM2 mutant ECs exhibit a competitive angiogenic advantage, allowing them to assert a stalk position upon neighboring wild-type ECs. Notably, CCM2 acts as a gatekeeper of ROCK-dependent mechanotransduction, attenuating forces on the ECM through both ROCKs and regulating matrix degradation through ROCK1. Our finding that modulation of ROCK-dependent force exertion also modifies the assignment of tip-stalk phenotypes represents, to our knowledge, the first evidence for a cell mechanics-driven mechanism for attracting wild type ECs in CCM and identifies dysregulated cell mechanics as a “third hit” for CCM pathogenesis.

**Figure 5.**
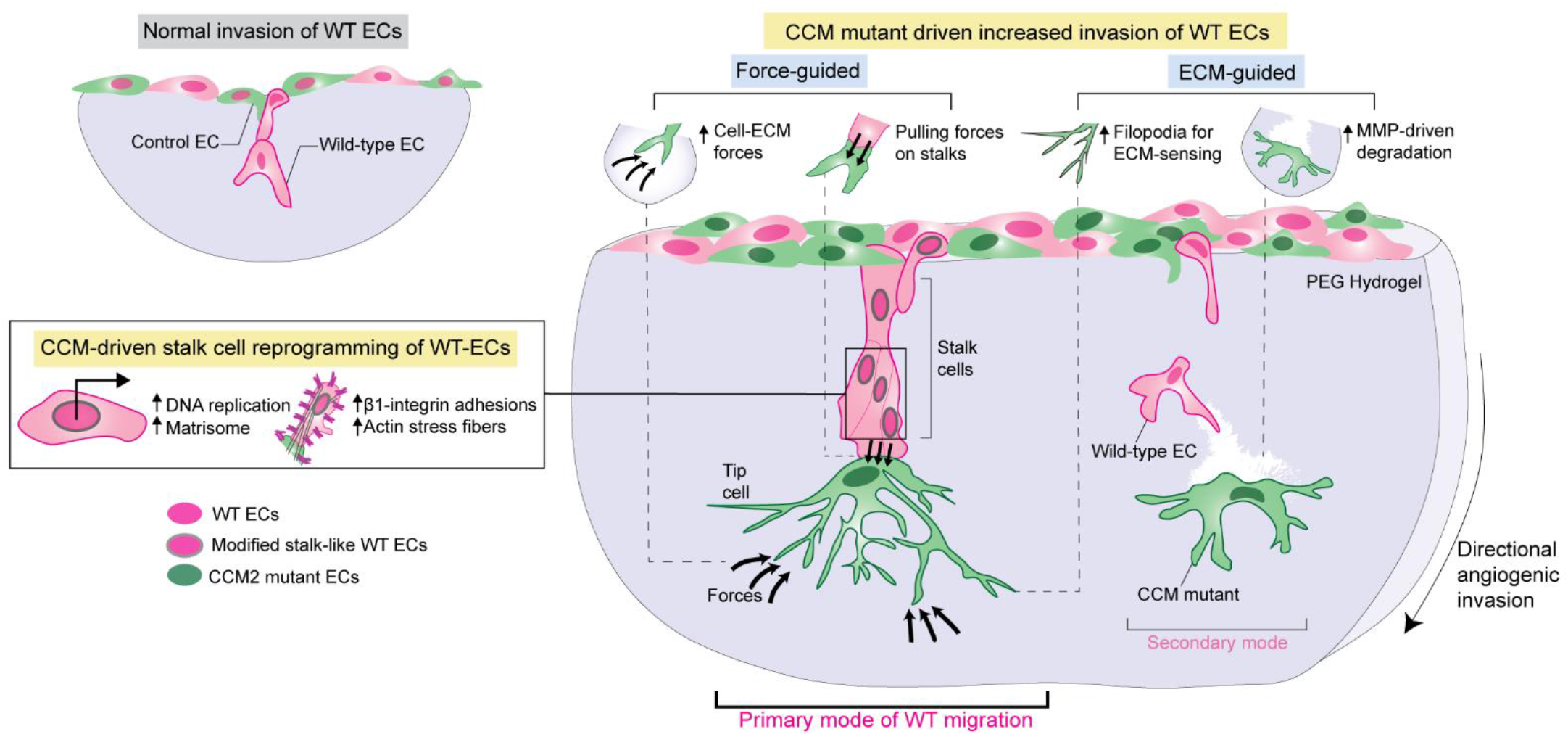
Proposed model for wild-type EC recruitment by siCCM2 mutant ECs. CCM2-silenced endothelial cells enhance angiogenic invasion of neighboring wild-type cells through force-guided and extracellular matrix-guided mechanisms. Mechanically hyperactive CCM2-silenced tips guide wild-type cells by exerting and transmitting pulling forces and by leaving degraded paths in the matrix as cues to further promote invasion. This further leads to a reinforcement of β1 integrin-dependent adhesions and actin stress fibers. This is accompanied by a transcriptomic reprogramming of wild-type ECs into stalk cells presenting increased proliferation.

Our findings demonstrate that the aggravation of force exertion on the ECM by CCM2 mutants can be felt by follower wild type ECs. While it is plausible that there is always force transmission between tip and stalk cells to sustain collective migration during physiological sprouting, the substantial increase in cell-ECM forces due to CCM2 mutants suggest that any force transmission between a mutant leader and a follower would likely be amplified, given also the force equilibrium boundary conditions. Unfortunately, quantifying and comparing this phenomenon in our model was impossible because wild type ECs rarely formed mosaic sprouts with control ECs, and they never occupied stalk positions behind them. This even suggested a complete lack of physical intercellular communication between populations in CT mosaics. Importantly, the proposed mechanism of hypercontractile mutants pulling forward passive wild type ECs was also associated with the reinforcement of the actin cytoskeleton and β1 focal adhesions in follower WT ECs. We speculate that a mechanical continuum is established between the cytoskeletal machinery of the mutants and wild-type ECs, although the underlying mechanisms remain to be addressed. It is conceivable that localized high forces at mosaic cell-cell junctions induce a force-dependent global activation of β1 integrins and actin assembly in WT ECs, similar to the mechanism driving the reinforcement of β1 integrin-dependent adhesive structures in response to PECAM mechanical activation^48^. Besides enabling focal adhesions and invasion, CCM2 mutants also initiate a transcriptomic reprogramming of wild type ECs into a stalk-like identity, characterized by the downregulation of tip cell markers and the upregulation of matrix deposition genes.

Cell proliferation and recruitment both play significant roles in the pathogenesis of CCM lesions. However, their respective contributions and temporal sequence remain unclear and may vary depending on the specific CCM gene involved. In zebrafish embryonic CCM1 knockout models and acute conditional CCM2 knockout mouse models induced shortly after birth, there is no evidence of EC proliferation^49,50^. In chronic CCM3 mouse models, an initial wave of proliferation of mutant cells is followed by the recruitment of wild-type ECs, which further promote the growth of multicavernous lesions^8,9^. Notably, in symptomatic human CCM1-3 lesions, wild-type ECs are present in significant numbers, supporting the idea that recruitment of wild-type ECs is a common mechanism in familial CCMs. Most remarkably, our study has shown that when exposed to a CCM2 mutant environment, wild-type ECs activate their DNA replication and proliferation programs, leading to increased DNA synthesis. To our knowledge, this is the first study suggesting an increase in proliferation of non-mutant endothelial cells in the context of CCMs. In contrast, CCM2 mutants exhibit cell cycle arrest associated with senescence^24^. Based on these results, we speculate that the proliferative response of wild-type cells might be the driver of multicavernous lesion growth after initial clonal expansion of CCM mutants. This model is supported by evidence that PCNA-positive proliferative ECs in human CCMs^51^ are likely present in zones of positive immunoreactivity for the corresponding CCM protein^52^. Further in vivo investigations will be required to ascertain this important change in paradigm in our understanding of CCM development.

Mechanistically, given the evidence of force transmission between mutant and WT cells, we speculate that the reprogramming of WT cells into a proliferative state may be mechanically driven. How mechanical signals from cell-cell forces are then transmitted to the nucleus to activate a reprogramming remains to be explored. Existing studies provide some essential clues; for instance, increased cell-matrix interactions and actin stress fibers in both cardiomyocytes and epithelial monolayers lead to YAP activation, driving increased proliferation^53,54^. The application of external stretch to polarize the collagen ECM in epithelial networks or pulling forces experienced from neighboring cells in MDCK monolayers, both stimulated cell proliferation through ERK activation, providing evidence of a direct force-based mechanism^55–57^.

Studies on CCM pathogenesis have proven exceptionally valuable in shedding light on vascular development and disease. Our unique perspective on the mechanics of the heterogenous vascular tissue during lesion growth opens up a complementary avenue for further in-vivo exploration. Our novel computational development to quantify cell-cell forces within a 3D multicellular system represents a significant advancement for other fields such as mathematical modeling of physiological processes^58^. Furthermore, our comprehensive methodology integrating 3D in-vitro systems, biophysical computations and molecular biology techniques holds broader application for exploring the role of mechanics and cellular interactions in the progression of other diseases characterized by aberrant angiogenesis.

## Supporting information

Extendend Data and Supplementary Information

## Methods

### Cell culture and siRNA silencing of HUVECs

Pooled Human Umbilical Vein Endothelial Cells (HUVEC) were obtained from Pelobiotech. HUVECs received at P0 were expanded over 2 passages in 100µg/ml collagen 1 (from rat tail, BD) coated flasks in complete EGM-2 medium (Lonza) supplemented with 100 U/ml penicillin and 100 μg/ml streptomycin at 37°C in a 5% CO2 - 3% O2 humidified chamber. HUVECs at passage 3 were transfected twice at 24 h-intervals with 30 nM siRNA and Lipofectamine RNAi max (Life Technologies, ref. 13778-150) according to the manufacturer’s instructions in OptiMEM at 37°C in a regular 5% CO_2_ in a humidified chamber. For double transfections, 30 nM of each siRNA duplexes (Dharmacon smartpool ON-TARGET plus Thermo Scientific) was used: Non-targeting siRNA #1 (ref. D-001810-01), CCM2 siRNA (ref. L-014728-01), ROCK1 siRNA (ref. L-ref. L-003536-00) and ROCK2 siRNA (ref. L-004610-00). siRNA transfected HUVECs were transduced with adenoviral LifeAct-GFP2 (Ibidi, Martinsried, Germany) at a multiplication of infection (MOI) of 5 and incubated for 16h while the wild-type HUVECs used in mosaics were similarly transduced with LifeAct-RFP2.

### PEG hydrogel and 3D invasion assay

A previously reported 3D angiogenesis assay^59^ was adapted to incorporate a polyethylene glycol based hydrogel (PEG) (Fig. 1a). Degradable PEG hydrogel precursors and buffers were prepared as previously described ^39,60^ and supplemented with 1:60 far-red fluorescent beads (FluoSpheres Carboxylate-Modified Microspheres, 200 nm, Invitrogen). Briefly, a PEG hydrogel was prepared by using activated FXIII to enzymatically crosslink an MMP-sensitive peptide modified PEG precursor (8-arm 40kDa), lysine-RGD peptide at 50 µM (Pepmic), and 0.1-0.5 µM Sphingosine-1-Phosphate (Sigma-Aldrich). Sonicated beads, milliQ water, 10X Buffer, lysine-RGD and S1P were first mixed in that order, followed by 10X FXIII which activated crosslinking. We used a wt/vol hydrogel composition of 1.2% at Young’s modulus (E) of 520Pa for 3D TFM experiments to obtain large enough matrix deformations, and 1.5% at E=800 Pa for all other experiments. The pre-polymerized PEG solution was pipetted into a pre-cut imaging chamber (Invitrogen Secure-Seal^TM^ S24732) adhered to the glass bottom of a 24 well plate (Cellvis, #P24-0-N), and allowed to crosslink at room temperature for 30 minutes. 40,000 HUVECs were seeded per chamber to obtain a confluent monolayer by incubating the dish vertically for 1 hour at 37°C, 5% CO_2_ leading to cell attachment onto the PEG meniscus. Dishes were then placed horizontally and incubated for 18 hours in complete growth medium. Samples were either imaged live for traction force microscopy experiments or fixed for 1.5h in 4% paraformaldehyde for immunostaining or invasion measurements,.

### Conditioned media experiments

MiliQ water normally used for PEG hydrogel preparation was replaced with 48 h conditioned media from control or CCM2 siRNA-transfected HUVECs. Lifeact-RFP labeled wild-type HUVECs were seeded in the invasion assay as described above and fixed after 18h of invasion and quantified (Extended Data Fig. 3-A, B).

### Image acquisition

For overnight timelapse imaging during sprouting, samples were placed at the microscope immediately after cell attachment and media addition. All images were acquired on Leica SP8 confocal microscopes with a 25x 0.95 NA water-immersion objective (for immunostaining), 20x 0.4 NA dry objective (for non-mosaic 3D TFM, overnight timelapse imaging), 10x 0.4 NA dry objective (for fixed invasion imaging) or on a Nikon confocal microscopy (A1R+MP) combined with an adaptive optics (AO) corrector at the MicroCell facility of IAB, Grenoble (for mosaic 3D TFM and a subset of overnight timelapse imaging).

### Hydrogel mechanical characterization

Homebuilt cylindrical molds treated with Sigmacote were used to prepare circular, disk-shaped hydrogels with a diameter of 7mm and a height of ∼2mm (Supplementary Fig. 1D). After crosslinking for 30 minutes at room temperature, hydrogels were carefully scooped using a stainless steel micro spoon spatula and transferred to phosphate-buffer saline (PBS)-filled petri dishes and maintained overnight at 4°C. A micro-laser scanning device (in house developed Micro Laser Scanner in collaboration with Acacia Technology, KU Leuven Core Facility for Biomechanical Experimentation – FIBEr) was used to measure the diameter and thickness of the sample before using an unconfined compression testing protocol^61^. This was crucial for applying a strain-controlled unconfined compression to the sample using a mechanical loading setup (Bose ElectroForce, TA Instrument Company). Our unconfined compression loading protocol comprised of a 20% applied strain at a rate of 1% per second adjusted to the measured height of each individual sample, followed by 15 minutes of relaxation^61,62^. The reaction force was recorded during the compression-relaxation experiment. Each PEG hydrogel with different concentration was measured from three independent trials in triplicates.

To obtain the mechanical properties of the hydrogels, an inverse finite element (FE)-based mechanical characterization approach was implemented. To this end, an FE simulation of the unconfined compression tests for each sample was created in Abaqus using a poro-hyperelastic material model as previously reported^62^. Briefly, the gels were simulated using 2D axisymmetric models and the material model consists of a Neo-Hookean solid matrix and a fluid phase. The biphasic behavior was simulated using Darcy’s law. Three material constants of the poro-hyperelastic material model include Young’s modulus (E) and Poisson’s ratio (υ) of the solid matrix and permeability parameter (k). An inverse parameter fitting algorithm was then used to fit the reaction force versus time obtained from the FE model to the experimental measurements by modifying the three material constants. For this purpose, *fminsearch* optimization function in MATLAB was used. The mechanical properties were determined for the three compression-relaxation testing trials of the PEG hydrogels and an average value was calculated for each material constant of each PEG hydrogel.

### Immunofluorescence staining

Sprout-laden PEG hydrogels were fixed for 2 h in 4% methanol-free paraformaldehyde (Thermo-Scientific) followed by washing with PBS. Samples were then treated with non-permeabilizing blocking solution constituting 1% BSA, 10% goal serum and 0.05% Tween 20 for 5h. Primary antibodies suspended in blocking solution were applied to fixed and blocked samples and left overnight at 4°C. Anti-activated β1 integrin clone 9EG7 (BD Biosciences #553715, 1:100) and recombinant anti-MMP14 antibody EP1264Y (abcam #ab51074, 1:100) were used as primary antibodies. Five PBS+Tween 20 washes were performed over 9 h followed by overnight immunolabeling with secondary antibodies diluted in non-permeabilizing blocking solution. Goat anti-rabbit or anti-rat IgG (H + L) highly cross-adsorbed secondary antibodies conjugated with Alexa Fluor AF 488, AF 546, AF 647 (Invitrogen, 1:500) and phalloidin conjugates with Atto 647 (Sigma-Aldrich, 1:1000) were used. Samples were then extensively washed over 24h with PBS + Tween 20 before imaging.

### EdU proliferation assay

Twenty-four hours after the second siRNA transfection, transfected HUVECs were combined with CellTracker green CMFDA dye (Invitrogen) labeled wild-type HUVECs in a 1:1 ratio and seeded at a final density of 7.5 X 10^4^ in 24-well plates coated with collagen 1 (from rat tail, BD). After culturing for 48h in complete medium at 37°C and 5% CO_2_, cells with active DNA synthesis were labeled using the edU Staining Proliferation Kit (iFluor 647, abcam) per manufacturer instructions.

### 3D traction force microscopy image acquisition

Confocal images were acquired at a 512x512 resolution using a 20x 0.4 NA dry objective at 2X zoom, a 56.6µm pinhole set to 1AU and bidirectional imaging with phase correction. The excitation wavelengths were 488nm (green cell channel), 552nm (red cell channel) and 638nm (far-red beads channel). The emission wavelengths for the PMT detectors were 498-538nm, 557-619nm and 673-704nm respectively. For mosaic experiments, the images were acquired sequentially to minimize crosstalk between the green and red channels, while for non-mosaic experiments the green cell channel and far-red beads channel were imaged simultaneously. The voxel size was set to 0.568×0.568×1 µm^3^ (see example images in Fig. 1f, top row). All TFM experiments were performed on Leica SP8 microscopes equipped with a temperature controlled stage maintained at 37°C and a humidified 5% CO_2_ environment. First, all positions of interest from each conditions in a well plate were identified and marked. The stressed (actively pulling) state was first acquired followed by a treatment with Cytochalasin D (dissolved in DMSO, Sigma Aldrich) at a final concentration of 4 µM for 1 hour. Finally, the relaxed (mechanically inactive) state was acquired in the same location.

### 3D traction force microscopy analysis

3D TFM data was analyzed using the TFMLAB software toolbox^63^. Briefly, bead and sprout images were filtered and enhanced. Sprout images were binarized to obtain a 3D cell mask. Any possible microscope stage drifts that occurred between the stressed and the relaxed states were corrected for using phase correlation. Free form deformation-based image registration was used to measure the 3D bead displacements^64^. After displacement measurement, regions of interest around the sprout tips were selected within TFMLAB. The tractions exerted by the cells were calculated in a Finite Element (FE) framework using our previously described physically based inverse method^65,66^. Briefly, this method searches for a displacement field that is as similar as possible to the measured one while satisfying the force equilibrium condition in the hydrogel domain. The hydrogel domain was modeled as a linear elastic material with E = 520 Pa (see hydrogel mechanical characterization section) and ν = 0.3^67^. TFMLAB assumes a negligible stiffness for the cell internal domain (Young’s modulus of 10^-6^ Pa). More details about the implementation of the method can be found in Supplementary note 1. We also developed a novel approach to quantify cell-cell forces from TFM data. A detailed description of this development can be found in Supplementary note 2. In Fig. 3c,d, we defined which sprouts were exerting forces by selecting the ones that generated matrix displacements larger than the noise level. Here, the noise level corresponds to matrix displacements greater than the pixel size, which is ∼0.5 μm.

### Quantification of filopodia and actin

3D fluorescence image stacks of phalloidin labeled sprouts were maximally projected and filtered with a median filter (function medfilt2 of MATLAB. Filtered images were binarized using a manually adjusted intensity threshold to provide a spout mask. The total actin intensity metric (Fig. 1c) was obtained by summing intensity values of the pixels within this mask and normalizing to the number of nuclei. The number of filopodia in sprout tips (<2 μm thick) were quantified by applying an opening operation on the binary mask using a disk-shaped structuring element of a 7 pixels radius. The resulting binary image was subtracted from the sprout mask and small objects (<50 pixels) were removed. The count of the connected components in the resultant image normalized for the number of nuclei provided the number of filopodia in Fig. 1d. The maximum filopodia length per sprout was measured manually using ImageJ. To quantify branchiness, the binary sprout masks were skeletonized followed by the removal of spurious structures shorter than 10μm (functions bwskel and bwmorph of MATLAB with options ‘MinBranchLength’, ‘spur’ and ‘hbreak’). Finally, the number of branch points (points where a skeleton branch ends) were counted in the resultant skeleton (Supplementary Fig. 1A depicts all operations).

### Quantification of matrix degradation

Gelatin labeled with Alexa 647 dye^68^ was combined at 2x final concentration in the pre-polymerization PEG solution to give a uniformly fluorescent hydrogel. Hydrogels were visualized after angiogenic invasion using a 647 laser to identify degraded (black zones) regions. The following image processing workflow was used to quantify the degraded area (Supplementary Figure 7B). Gelatin-Alexa647 images were filtered with a disk-shaped top-hat filter to remove uneven background illumination, followed by a smoothing Gaussian filter and a contrast stretching operation. A minimum-intensity projection was then applied to obtain the complementary image. The degraded region was binarized using a manually adjusted intensity threshold and the degradation area was computed around the sprout of interest, normalized to the number of nuclei, and displayed in Fig. 1j and Extended Data Fig. 4.

### Quantification of MT1-MMP, and β1 integrin

MT1-MMP and β1-integrin immunofluorescence were quantified based on 3D confocal stacks images. Images were first maximally projected followed by a 5x5 wiener filter to reduce background noise and a contrast stretching operation. All images were processed equally such that the quantifications are comparable. The intensity values of the pixels within a cropped region of interest were summed, normalized by the number of nuclei, and displayed in Fig. 1l and Fig. 3j.

### Quantification of invasion distance

100μm stacks of 3D confocal images were maximally projected using ImageJ (Version 1.52a) and the maximum invasion distance was manually measured over ∼10 field of views (FOVs) of the monolayer for each replicate at multiple positions within each FOV. The mean invasion distance for each condition quantified per trial was then normalized to the control of that trial (Fig. 2b). This was done for the cell population of interest by splitting channels based on fluorescence.

### Quantification of dynamic invasion by cell tracking

Confocal images stacks (100μm thick, 1 μm step size) of mosaic sprouting acquired every 45 minutes over 16 hours were maximally projected in ImageJ and converted to .avi files in RGD format. First, an automated algorithm was developed to automatically segment the hydrogel domain (Supplementary Fig. 1C). Specifically, the entire time-lapse of both cell channels was first maximum-intensity projected). The resultant image was then filtered, its complementary obtained and binarized. Only the largest component of the image corresponding to the hydrogel domain was kept. This was followed by obtaining the convex hull of the binary image (MATLAB function bwconvhull) and calculating its distance transform.

Each image of the timelapse was then binarized, and the inner holes were removed (Supplementary Fig. 1C). Spurious binary objects with areas lower than ∼30µm^3^ or those that did not coincide with the segmented hydrogel domain were removed. Binarized cells in the last time point were tracked by comparing the distances between their centroids and those of the binarized cells of the previous time point. The nearest neighbor was selected as the match, consequently attributing to the pertinent label. This process was repeated iteratively until covering the entire timelapse. Notably, in instances where a cell division event occurred, the existing label was allocated to one of the cells while a new label was assigned to the other cell. The invasion distance of each cell was tracked by storing the maximum value of the distance transform of the hydrogel domain within the cell’s binary mask. The results are shown in Fig. 2d and Supplementary Video 4. Modes of invasion of wild-type ECs shown in Fig. 2f, g were quantified by manually counting the number of WT ECs in each mode at the first moment of invading the hydrogel as visualized in overnight timelapses.

### Quantification of EdU staining

The WT cell, nuclei, and EdU-stained images underwent a series of preprocessing steps programmed in MATLAB to enhance their quality. Initially, denoising was performed by applying a median filter, followed by a Gaussian filter, and a contrast stretching operation using the respective functions medfilt2, imgaussfilt, and imadjust. Subsequently, a binarization process was applied and any holes within these binary images were filled using the imbinarize and imfill functions. To address cases where nuclei were in close proximity or overlapping due to segmentation inaccuracies, we employed the watershed algorithm on the binary nuclei image, which separated closely situated nuclei watershed function. We then counted the EdU nuclei that were overlapping with both the WT cell and nuclei binary masks. We reported the percentage of EdU positive WT cells in Fig. 4k.

### Single-cell RNA sequencing sample preparation

Three independent trials of mosaic invasion samples were prepared for control (WT-RFP + siCT-GFP ECs) and CCM2^KD^ (WT-RFP + siCCM2-GFP ECs) conditions and allowed to sprout for 24h into degradable PEG 1.5% hydrogels with a high concentration of S1P (400nM) to allow for significant hydrogel invasion by both cell populations. Hydrogels were then collected using a spatula into a falcon tube and dissociated into single cells using an incubation period of 15 min in trypsin, thereby leaving behind the cells attached in 2D on the dish bottom. The solutions were neutralized with a 1:10 volume of EGM-2 without antibiotics and centrifuged for 5 min at 300 RCF, followed by 10 min at 500 RCF. After decanting the supernatant, the cell pellets were resuspended in 200 μL of EGM-2 and put on ice. Cells were isolated by fluorescence activated cell sorting (FACS) by flow cytometry on a Aria IIu (BD Biosciences). Discrimination of cell populations was based on fluorescence at 530/30 nm from 488 nm excitation and 610/20 nm from 561 nm excitation to identify respectively cells expressing Lifeact-GFP or Lifeact-RFP. Cells were run through a 100 um nozzle under 20 psi at a flow rate of 1 and recovered simultaneously in tubes in a 4-way purity mode. The sorted cells were labeled as WT, WT-CCM2, siCCM2 and frozen in EGM-2 + 40% FCS + 10% DMSO.

After thawing the cells, their viability (>80%) and aggregation were tested before starting the single-cell library preparation. The concentration of freshly dissociated cells was adjusted to 1000 cells/μL in HBSS/0.05% BSA, and then the 10x Chromium Controller and the Chromium Single Cell 3′ Reagent kit V3.1 were used to perform the scRNAseq experiments. Library construction was performed according to the manufacturer’s instructions using the Chromium Chip B Single Cell kit and Chromium Multiplex Kit (10X Genomics, Pleasanton, CA, USA). Sequencing was performed in paired-end mode with HighOutput flow cell (read 1: 28 cycles and read 2: 90 cycles) and a NextSeq 550 sequencer (Illumina, San Diego, CA, USA) at the Transcriptome Core Facility of the Institute in Regenerative Medicine and Biotherapy, CHU-Inserm-UM Montpellier, France. CellRanger was used to process scRNAseq data. We first use CellRanger mkfastq to convert and demultiplex the bcl files (from illumina sequencer, raw data) in fastq files for each sample. Then we use CellRanger count to generate the matrix data containing gene counts for each cell per sample. Fastq files were aligned to the human genome reference sequence GRCh38.

### Single-cell sequencing data processing and clustering

After QC steps we have retained a total of 1434: 464, 461 and 509 for the WT, WT-CCM2 and CCM2^KD^ conditions respectively. We implemented the Seurat^69^ 4.3.0 pipeline to perform scRNAseq analysis. The data was normalized and 2000 highly variable genes (features) were identified using the FindVariableFeatures. The data was scaled and described using principal component analysis (PCA) using the RunPCA function. The data was clustered by the top 7 PCs and the FindNeighbors function parameter k.param = 5. The FindCluster resolution was set to 0.15. Data visualization was done using the Uniform Manifold Approximation and Projection (UMAP) dimensionality reduction technique using the RunUMAP package with the top 7 PCs and identified 7 clusters in the data. Using the top 50 upregulated genes we annotated the 7 clusters at a high-level as 1) Proliferation, 2) Quiescence, 3-5) Angiogenesis 1/2/3, 6) Vasculature remodeling, 7) RNA splicing. Genesets were created based on previous studies^69–78^ and geneset scores per data type were reported in Figure 4d,h and j. Normalized expression levels of *CD34* in *CD34*^+^ cells were extracted using the FetchData function. Average gene expressions were obtained by the AverageExpression function.

### Statistical analysis

Our data across all conditions to be compared did not pass the normality test. Hence, we employed an unpaired Mann Whitney non-parametric t-test and a Kruskal Wallis non-parametric ANOVA statistical test with corrections were appropriate with a 95% confidence interval (GraphPad Prism 9, Version 9.5.1, GraphPad Software, Inc.). Error bars always represent SEM. Statistical significance was considered for all comparisons with the following p-values <0.05(*) <0.01(**) <0.001(***). When determining significance difference between conditions, the multiple comparisons were performed comparing all conditions to the control unless otherwise specifically depicted. Measurements were taken from several distinct cell sprouts within a sample and the same cell sprout was repeatedly measured only for overnight imaging to study the dynamics.

## Data availability statement

Single-cell RNA sequencing row datasets generated in this study have been deposited in the Gene Expression Omnibus repository. Reviewer tokens are available upon request from A.S. All other data that support the findings of this study are available from the corresponding authors upon reasonable request.

## Code availability statement

TFMLAB code used to analyze 3D traction force microscopy data is open-source and can be accessed from the Gitlab repository: https://gitlab.kuleuven.be/MAtrix/Jorge/tfmlab_public. Implemented cell-cell force recovery and image analyses routines developed for all quantifications are available upon reasonable request.

## Acknowledgements

We thank the members of the Van Oosterwyck laboratory for thoughtful feedback; Transcriptom Core Facility of the Institute in Regenerative Medicine and Biotherapy, CHU-INSERM-UM Montpellier, https://irmb-montpellier.fr/ for the single-cell RNA Sequencing processing; and the MicroCell microscopy facility at IAB, R. Nuyts and A. Grichine for support with microscopy. These studies were supported by Research Foundation Flanders (FWO) PhD fellowship 1S68820N, postdoctoral fellowship 1259223N, projects G087018N, G0C2422N; infrastructure grant I009718N, KU Leuven grants IDN/19/031, iBOF/21/083 C; ERANET NEURON JTC 2022 call – project G0L1522N, Project PID2021-126051OB-C42 Ministerio de Ciencia e Innovación (MCI), Agencia Estatal de Investigación (AEI) and Fondo Europeo de Desarrollo Regional (FEDER), Marie Skłodowska–Curie Individual Fellowship (CREATION project: MSCA-IF-2019-893771).

## Author contributions

A.S. designed, conducted, imaged, analyzed, interpreted all experiments, and wrote the manuscript. J.B.F developed all image analysis tools, J.B.F and J.A.S developed the cell-cell force recovery algorithms and interpreted the results. M.P. performed FACS sorting of mosaic cell populations, S.A. generated and A.A. analyzed and interpreted the scRNAseq data. A.S. and E.D.V. conducted and analyzed experiments for hydrogel mechanical characterization while S.A.E. implemented the finite element analyses. J.D.J. conducted tissue culture experiments. A.S., J.B.F., A.A., J.D.J., J.A.S, M.P, S.A, E.D.V, S.A, S.A.E, A.R., E.F., and H.V.O. edited the manuscript. E.F. designed and interpreted experiments and wrote the manuscript. H.V.O. interpreted experiments and wrote the manuscript.

## Additional information

Extended data figures and supplementary information is available for this paper as a separate document.

## References

1. Barrasa-Ramos, S., Dessalles, C. A., Hautefeuille, M. & Barakat, A. I. Mechanical regulation of the early stages of angiogenesis. J R Soc Interface 19, (2022).

2. Zanotelli, M. R. & Reinhart-King, C. A. Mechanical forces in tumor angiogenesis. Adv Exp Med Biol 1092, 91–112 (2018).

3. Flournoy, J., Ashkanani, S. & Chen, Y. Mechanical regulation of signal transduction in angiogenesis. Front Cell Dev Biol 10, 933474 (2022).

4. Wang, D., Brady, T., Santhanam, L. & Gerecht, S. nature cardiovascular research The extracellular matrix mechanics in the vasculature. Nature Cardiovascular Research 2, 718–732 (2023).

5. Jung, K. H. et al. Cerebral Cavernous Malformations With Dynamic and Progressive Course: Correlation Study With Vascular Endothelial Growth Factor. Arch Neurol 60, 1613–1618 (2003).

6. Awad, I. A. & Polster, S. P. Cavernous angiomas: deconstructing a neurosurgical disease: JNSPG 75th Anniversary Invited Review Article. J Neurosurg 131, 1–13 (2019).

7. Snellings, D. A. et al. Cerebral Cavernous Malformation: From Mechanism to Therapy. Circ Res 129, 195–215 (2021).

8. Malinverno, M. et al. Endothelial cell clonal expansion in the development of cerebral cavernous malformations. Nat Commun 10, 2761 (2019).

9. Detter, M. R., Snellings, D. A. & Marchuk, D. A. Cerebral cavernous malformations develop through clonal expansion of mutant endothelial cells. Circ Res 123, 1143–1151 (2018).

10. Detter, M. R. et al. Novel Murine Models of Cerebral Cavernous Malformations. Angiogenesis 23, 651–666 (2020).

11. Rath, M., Pagenstecher, A., Hoischen, A. & Felbor, U. Postzygotic mosaicism in cerebral cavernous malformation. J Med Genet 57, 212–216 (2020).

12. Gault, J. et al. Cerebral Cavernous Malformations: Somatic Mutations in Vascular Endothelial Cells. Neurosurgery 65, 138 (2009).

13. Gault, J., Shenkar, R., Recksiek, P. & Awad, I. A. Biallelic Somatic and Germ Line CCM1 Truncating Mutations in a Cerebral Cavernous Malformation Lesion. Stroke 36, 872–874 (2005).

14. Akers, A. L., Johnson, E., Steinberg, G. K., Zabramski, J. M. & Marchuk, D. A. Biallelic somatic and germline mutations in cerebral cavernous malformations (CCMs): evidence for a two-hit mechanism of CCM pathogenesis. Hum Mol Genet 18, 919–930 (2009).

15. Whitehead, K. J. et al. The cerebral cavernous malformation signaling pathway promotes vascular integrity via Rho GTPases. Nature Medicine 2009 15:2 15, 177–184 (2009).

16. Borikova, A. L. et al. Rho kinase inhibition rescues the endothelial cell cerebral cavernous malformation phenotype. Journal of Biological Chemistry 285, 11760–11764 (2010).

17. Stockton, R. A., Shenkar, R., Awad, I. A. & Ginsberg, M. H. Cerebral cavernous malformations proteins inhibit Rho kinase to stabilize vascular integrity. Journal of Experimental Medicine 207, 881–896 (2010).

18. McDonald, D. A. et al. Fasudil decreases lesion burden in a murine model of cerebral cavernous malformation disease. Stroke 43, 571–574 (2012).

19. Shenkar, R. et al. Rho kinase inhibition blunts lesion development and hemorrhage in murine models of aggressive Pdcd10/Ccm3 disease. Stroke 50, 738–744 (2019).

20. Ren, A. A. et al. PIK3CA and CCM mutations fuel cavernomas through a cancer-like mechanism. Nature 2021 594:7862 594, 271–276 (2021).

21. Rath, M. et al. Contact-dependent signaling triggers tumor-like proliferation of CCM3 knockout endothelial cells in co-culture with wild-type cells. Cellular and Molecular Life Sciences 2022 79:6 79, 1–20 (2022).

22. Labernadie, A. et al. A mechanically active heterotypic E-cadherin/N-cadherin adhesion enables fibroblasts to drive cancer cell invasion. Nature Cell Biology 2017 19:3 19, 224–237 (2017).

23. Baschieri, F. et al. Fibroblasts generate topographical cues that steer cancer cell migration. Sci Adv 9, (2023).

24. Vannier, D. R. et al. CCM2-deficient endothelial cells undergo a ROCK-dependent reprogramming into senescence-associated secretory phenotype. Angiogenesis 24, 843–860 (2021).

25. De Smet, F., Segura, I., De Bock, K., Hohensinner, P. J. & Carmeliet, P. Mechanisms of Vessel Branching. Arterioscler Thromb Vasc Biol 29, 639–649 (2009).

26. Fischer, R. S., Lam, P. Y., Huttenlocher, A. & Waterman, C. M. Filopodia and focal adhesions: An integrated system driving branching morphogenesis in neuronal pathfinding and angiogenesis. Dev Biol 451, 86–95 (2019).

27. Lisowska, J. et al. The CCM1-CCM2 complex controls complementary functions of ROCK1 and ROCK2 that are required for endothelial integrity. J Cell Sci 131, jcs.216093 (2018).

28. Newell-Litwa, K. A. et al. ROCK1 and 2 differentially regulate actomyosin organization to drive cell and synaptic polarity. Journal of Cell Biology 210, 225–242 (2015).

29. Vaeyens, M.-M. et al. Matrix deformations around angiogenic sprouts correlate to sprout dynamics and suggest pulling activity · In vitro model · Endothelial invasion · Extracellular matrix · Collagen · Cytoskeleton · Mechanobiology · Mechanotransduction · Cellular forces · Pulling forces · Computer model · Image processing · Confocal microscopy. Angiogenesis 23, 315–324 (123AD).

30. Kretschmer, M., Rüdiger, D. & Zahler, S. Mechanical Aspects of Angiogenesis. Cancers 2021, Vol. 13, Page 4987 13, 4987 (2021).

31. Infante, E. et al. LINC complex-Lis1 interplay controls MT1-MMP matrix digest-on-demand response for confined tumor cell migration. Nature Communications 2018 9:1 9, 1–13 (2018).

32. Wang, W. Y., Jarman, E. H., Lin, D. & Baker, B. M. Dynamic Endothelial Stalk Cell–Matrix Interactions Regulate Angiogenic Sprout Diameter. Front Bioeng Biotechnol 9, 187 (2021).

33. Schmitt, C. A., Wang, B. & Demaria, M. Senescence and cancer — role and therapeutic opportunities. doi:10.1038/s41571-022-00668-4.

34. Yoon, C. et al. Myosin IIA–mediated forces regulate multicellular integrity during vascular sprouting. Mol Biol Cell 30, 1974 (2019).

35. Maruthamuthu, V., Sabass, B., Schwarz, U. S. & Gardel, M. L. Cell-ECM traction force modulates endogenous tension at cell-cell contacts. Proc Natl Acad Sci U S A 108, 4708–4713 (2011).

36. Faurobert, E. et al. CCM1-ICAP-1 complex controls ??1 integrin-dependent endothelial contractility and fibronectin remodeling. Journal of Cell Biology 202, 545–561 (2013).

37. Engler, A. J., Sen, S., Sweeney, H. L. & Discher, D. E. Matrix Elasticity Directs Stem Cell Lineage Specification. Cell 126, 677–689 (2006).

38. Carley, E., King, M. C. & Guo, S. Integrating mechanical signals into cellular identity Cell Biology. Trends Cell Biol 32, (2022).

39. Abdel Fattah, A. R., et al. Actuation enhances patterning in human neural tube organoids. Nature Communications 2021 12:1 12, 1–13 (2021).

40. Barriga, E. H., Franze, K., Charras, G. & Mayor, R. Tissue stiffening coordinates morphogenesis by triggering collective cell migration in vivo. Nature 554, (2018).

41. Ren, J. et al. Somatic variants of MAP3K3 are sufficient to cause cerebral and spinal cord cavernous malformations. Brain 146, 3634–3647 (2023).

42. Siemerink, M. J. et al. CD34 marks angiogenic tip cells in human vascular endothelial cell cultures. Angiogenesis 15, 151 (2012).

43. Langlois, B. et al. AngioMatrix, a signature of the tumor angiogenic switch-specific matrisome, correlates with poor prognosis for glioma and colorectal cancer patients. Oncotarget 5, 10529 (2014).

44. Sung, J. Y. & Cheong, J. H. The Matrisome is Associated with Metabolic Reprograming in Stem-Like Phenotypes of Gastric Cancer. Cancers (Basel) 14, (2022).

45. Pietilä, E. A. et al. Co-evolution of matrisome and adaptive adhesion dynamics drives ovarian cancer chemoresistance. Nat Commun 12, (2021).

46. Abdelilah-Seyfried, S., Tournier-Lasserve, E. & Derry, W. B. Blocking Signalopathic Events to Treat Cerebral Cavernous Malformations. Trends Mol Med 26, 874–887 (2020).

47. Gaggioli, C. et al. Fibroblast-led collective invasion of carcinoma cells with differing roles for RhoGTPases in leading and following cells. Nature Cell Biology 2007 9:12 9, 1392–1400 (2007).

48. Collins, C. et al. Localized Tensional Forces on PECAM-1 Elicit a Global Mechanotransduction Response via the Integrin-RhoA Pathway. Current Biology 22, 2087–2094 (2012).

49. Hogan, B. M., Bussmann, J., Wolburg, H. & Schulte-Merker, S. ccm1 cell autonomously regulates endothelial cellular morphogenesis and vascular tubulogenesis in zebrafish. Hum Mol Genet 17, 2424–2432 (2008).

50. Boulday, G. et al. Developmental timing of CCM2 loss influences cerebral cavernous malformations in mice. Journal of Experimental Medicine 208, 1835–1847 (2011).

51. Notelet, L., Houtteville, J. P., Khoury, S., Lechevalier, B. & Chapon, F. Proliferating cell nuclear antigen (PCNA) in cerebral cavernomas: an immunocytochemical study of 42 cases. Surg Neurol 47, 364–370 (1997).

52. Wouters, V. et al. Hereditary cutaneomucosal venous malformations are caused by TIE2 mutations with widely variable hyper-phosphorylating effects. European Journal of Human Genetics 2010 18:4 18, 414–420 (2009).

53. Li, X., McLain, C., Samuel, M. S., Olson, M. F. & Radice, G. L. Actomyosin-mediated cellular tension promotes Yap nuclear translocation and myocardial proliferation through α5 integrin signaling. Development (Cambridge) 150, (2023).

54. Balasubramaniam, L. et al. Investigating the nature of active forces in tissues reveals how contractile cells can form extensile monolayers. Nature Materials 2021 20:8 20, 1156–1166 (2021).

55. Katsuno-Kambe, H. et al. Collagen polarization promotes epithelial elongation by stimulating locoregional cell proliferation. Elife 10, (2021).

56. Hino, N. et al. ERK-Mediated Mechanochemical Waves Direct Collective Cell Polarization. Dev Cell 53, (2020).

57. Hirashima, T., Hino, N., Aoki, K. & Matsuda, M. Stretching the limits of extracellular signal-related kinase (ERK) signaling-Cell mechanosensing to ERK activation. Curr Opin Cell Biol 2023, 102217 (2023).

58. Santos-Oliveira, P. et al. The Force at the Tip - Modelling Tension and Proliferation in Sprouting Angiogenesis. PLoS Comput Biol 11, 1004436 (2015).

## Methods References

59. Vaeyens, M. M. et al. Matrix deformations around angiogenic sprouts correlate to sprout dynamics and suggest pulling activity. Angiogenesis 23, 315–324 (2020).

60. Ranga, A. et al. 3D niche microarrays for systems-level analyses of cell fate. Nat Commun 5, 1–10 (2014).

61. Elahi, S. A. et al. Unconfined Compression Experimental Protocol for Cartilage Explants and Hydrogel Constructs: From Sample Preparation to Mechanical Characterization. Methods in Molecular Biology 2598, 271–287 (2023).

62. Elahi, S. A. et al. Guide to mechanical characterization of articular cartilage and hydrogel constructs based on a systematic in silico parameter sensitivity analysis. Journal of the Mechanical Behavior of Biomedical Materials 124, 104795 (2021).

63. Barrasa-Fano, J. et al. TFMLAB: A MATLAB toolbox for 4D traction force microscopy. SoftwareX 15, 100723 (2021).

64. Jorge-Peñas, A. et al. Free form deformation-based image registration improves accuracy of traction force microscopy. PLoS One 10, 1–22 (2015).

65. Sanz-Herrera, J. A., Barrasa-Fano, J., Cóndor, M. & Van Oosterwyck, H. Inverse method based on 3D nonlinear physically constrained minimisation in the framework of traction force microscopy. Soft Matter 17, 10210– 10222 (2021).

66. Barrasa-Fano, J. et al. Advanced in silico validation framework for three-dimensional traction force microscopy and application to an in vitro model of sprouting angiogenesis. Acta Biomater 126, 326–338 (2021).

67. Cappello, J., d’Herbemont, V., Lindner, A. & Roure, O. du. Microfluidic in-situ measurement of poisson’s ratio of hydrogels. Micromachines (Basel) 11, 318 (2020).

68. Sharma, V. P., Entenberg, D. & Condeelis, J. High-resolution live-cell imaging and time-lapse microscopy of invadopodium dynamics and tracking analysis. Methods in Molecular Biology 1046, 343–357 (2013).

69. Satija, R., Farrell, J. A., Gennert, D., Schier, A. F. & Regev, A. Spatial reconstruction of single-cell gene expression data. Nature Biotechnology 2015 33:5 33, 495–502 (2015).

70. Berenjeno, I. M., Núñez, F. & Bustelo, X. R. Transcriptomal profiling of the cellular transformation induced by Rho subfamily GTPases. Oncogene 26, (2007).

71. Fridman, A. L. & Tainsky, M. A. Critical pathways in cellular senescence and immortalization revealed by gene expression profiling. Oncogene 2008 27:46 27, 5975–5987 (2008).

72. Hernandez-Segura, A. et al. Unmasking Transcriptional Heterogeneity in Senescent Cells. Current Biology 27, (2017).

73. Schaefer, C. F. et al. PID: the Pathway Interaction Database. Nucleic Acids Res 37, D674–D679 (2009).

74. Subramanian, A. et al. Gene set enrichment analysis: A knowledge-based approach for interpreting genome-wide expression profiles. Proc Natl Acad Sci U S A 102, 15545–15550 (2005).

75. Liberzon, A. et al. The Molecular Signatures Database Hallmark Gene Set Collection. Cell Syst 1, (2015).

76. Naba, A., et al. The Matrisome: In Silico Definition and In Vivo Characterization by Proteomics of Normal and Tumor Extracellular Matrices. Mol Cell Proteomics 11, (2012).

77. Kanehisa, M., Furumichi, M., Sato, Y., Kawashima, M. & Ishiguro-Watanabe, M. KEGG for taxonomy-based analysis of pathways and genomes. Nucleic Acids Res 51, D587–D592 (2023).

78. Kanehisa, M. & Goto, S. KEGG: kyoto encyclopedia of genes and genomes. Nucleic Acids Res 28, 27–30 (2000).

