## Supplementary material for "Force-mediated recruitment and reprogramming of healthy endothelial cells drive vascular lesion growth": Extendend Data and Supplementary Information

### Additional Information

**This file includes:**

### Extended Data Figures

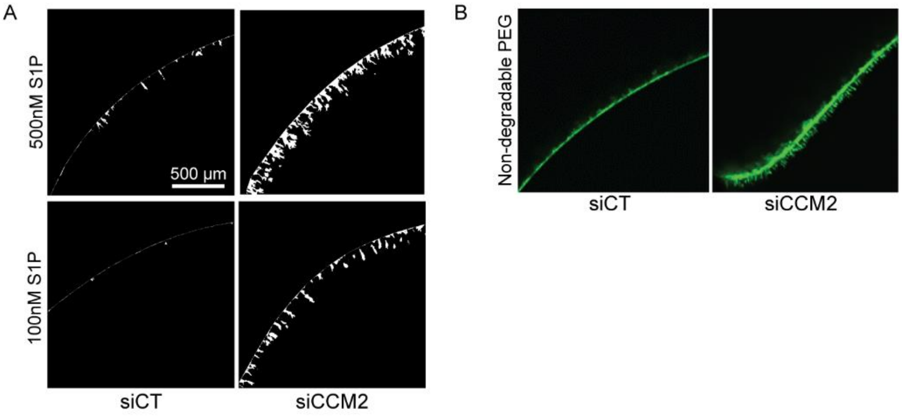

**Extended Data Fig. 1: siCCM2 ECs show increased ECM sensing.** Maximally projected sprouting images of siCT and siCCM2 EC monolayers into **A.** 1.5% PEG hydrogels with a low (100 nM) or high (500 nM) concentration of S1P after binarization, and **B.** a non-degradable 1.5% PEG hydrogel with 500 nM S1P.

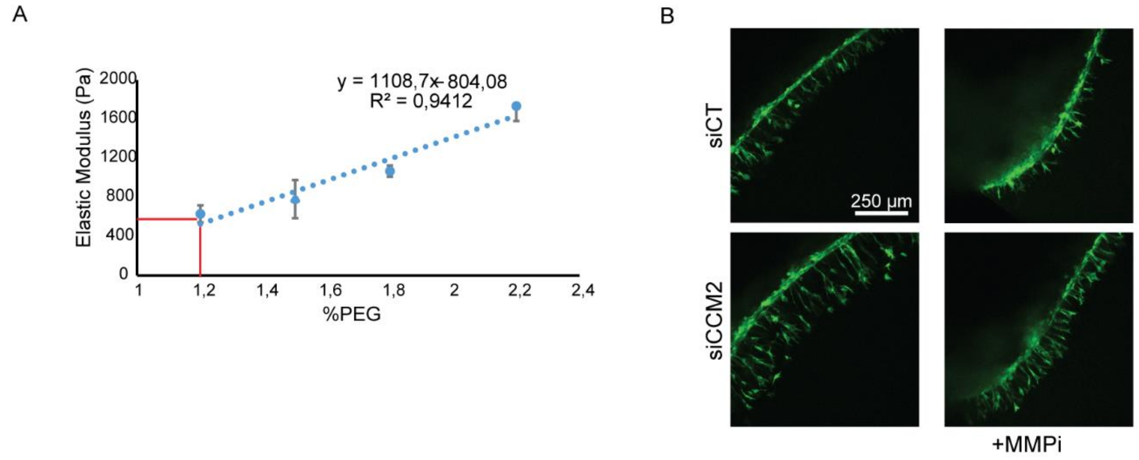

**Extended Data Fig. 2: 3D Traction force microscopy and degradation inhibition for siCCM2 ECs. A.** Elastic (Young's) modulus versus PEG percentage linear curve obtained from unconfined compression testing of degradable hydrogels embedded with beads. Data represents at least 3 independent trials. **B.** Maximally projected confocal sprouting images of siCT and siCCM2 EC monolayers with or without treatment with 25 µM MMPi.

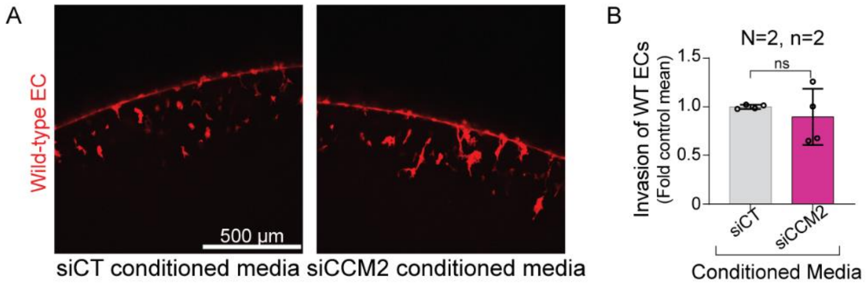

**Extended Data Fig. 3: Paracrine effects on WT ECs in 3D PEG invasion assay. A.** Maximally projected confocal images of wild-type ECs sprouting in PEG hydrogels prepared using media conditioned by siCT or siCCM2 ECs as the aqueous solution. **B.** Manual quantification of maximum invasion distance computed for two independent trials and ~10 images per trial and normalized to the control mean.

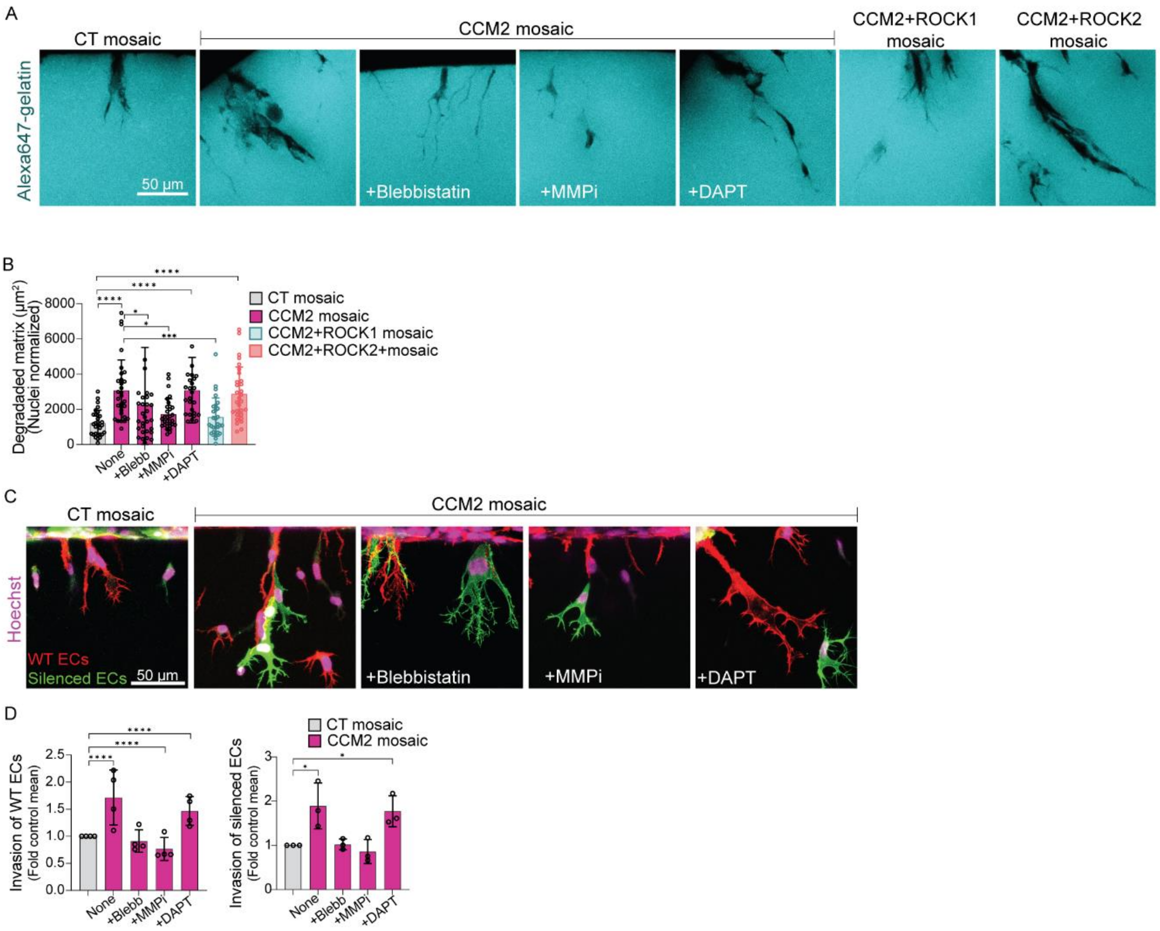

**Extended Data Fig. 4: ROCK-dependent myosin contractility and matrix degradation underlie enhancement of WT EC invasion by CCM2 ECs. A**. Representative minimal intensity projected confocal images of degradation of fluorescently labeled gelatin by CCM2 mosaics during overnight invasion with or without inhibition of contractility (+10µM Blebbistatin), degradation (+25 µM MMPi), Notch inhibition (+20µM DAPT), and by mosaics with CCM2 ECs additionally silenced for ROCK1 or ROCK2. **B**. Automated quantification of total degraded area normalized to the number of nuclei ($\mu m^2$, n=3, 25-30 sprouts/condition). **C.** Representative maximally projected confocal images of siCCM2 mosaic sprouting (1:1 WT (red) and silenced (green) ECs) with or without overnight treatment with blebbistatin, GM6001 or DAPT **D.** Quantifications of invasion distance of WT ECs and silenced ECs for treated and untreated mosaics (n=3, 25-30 positions/condition, quantified as averages after normalizing to the control for each trial).

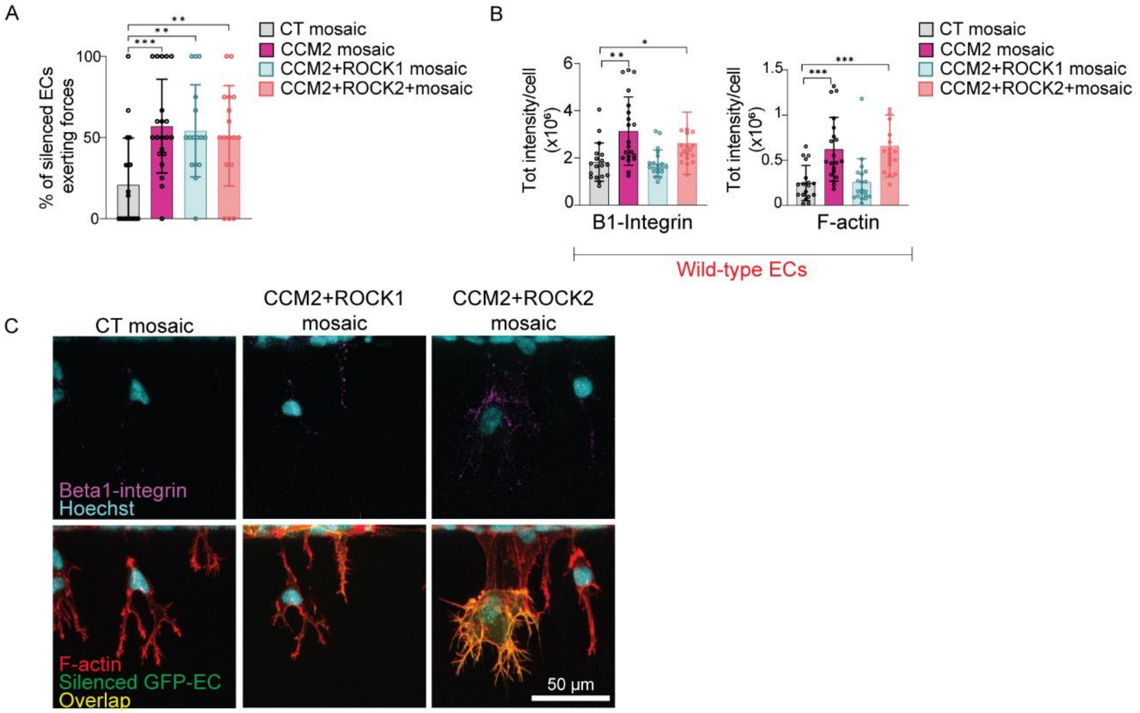

**Extended Data Fig. 5: ROCK-dependent modified mechanical behavior of WT ECs in CCM2 mosaics.** **A.** Quantification of the percentage of silenced ECs (in green) in CT, CCM2, CCM2+ROCK1 or CCM2+ROCK2 mosaics actively exerting forces on the matrix based on criteria in Fig. 3c (3 independent trials; CT, CCM2 mosaics, n=19; CCM2+ROCK1, CCM2+ROCK2 mosaics, n=21). **B.** Automated total intensity quantification per cell of β1 integrin expression and F-actin measured in regions containing only wild type ECs cropped from maximum projection images and normalized to the number of nuclei (2 independent trials; CT mosaic, n=18; CCM2, CCM2+ROCK1 mosaics, n=20; CCM2+ROCK2 mosaics, n=18). **C.** Representative confocal immunofluorescence images (maximum projections) of (**top**) β1 integrin (pink) and Hoechst-labeled nuclei (cyan), (**bottom**) fluorescently labeled silenced EC (green), F-actin (red) and the overlap (yellow). Cells that appear yellow are siCT, siCCM2+siR1 or siCCM2+siR2 ECs.

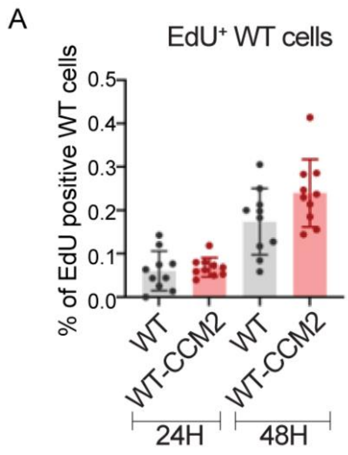

**Extended Data Fig. 6: EdU staining of WT ECs. A.** Percentage of EdU positive wild-type cells quantified for 1:1 mosaics of WT and siCT or siCCM2 cells culture for 24 or 48h. One independent trial with 10 images per dish.

### Supplementary Figures

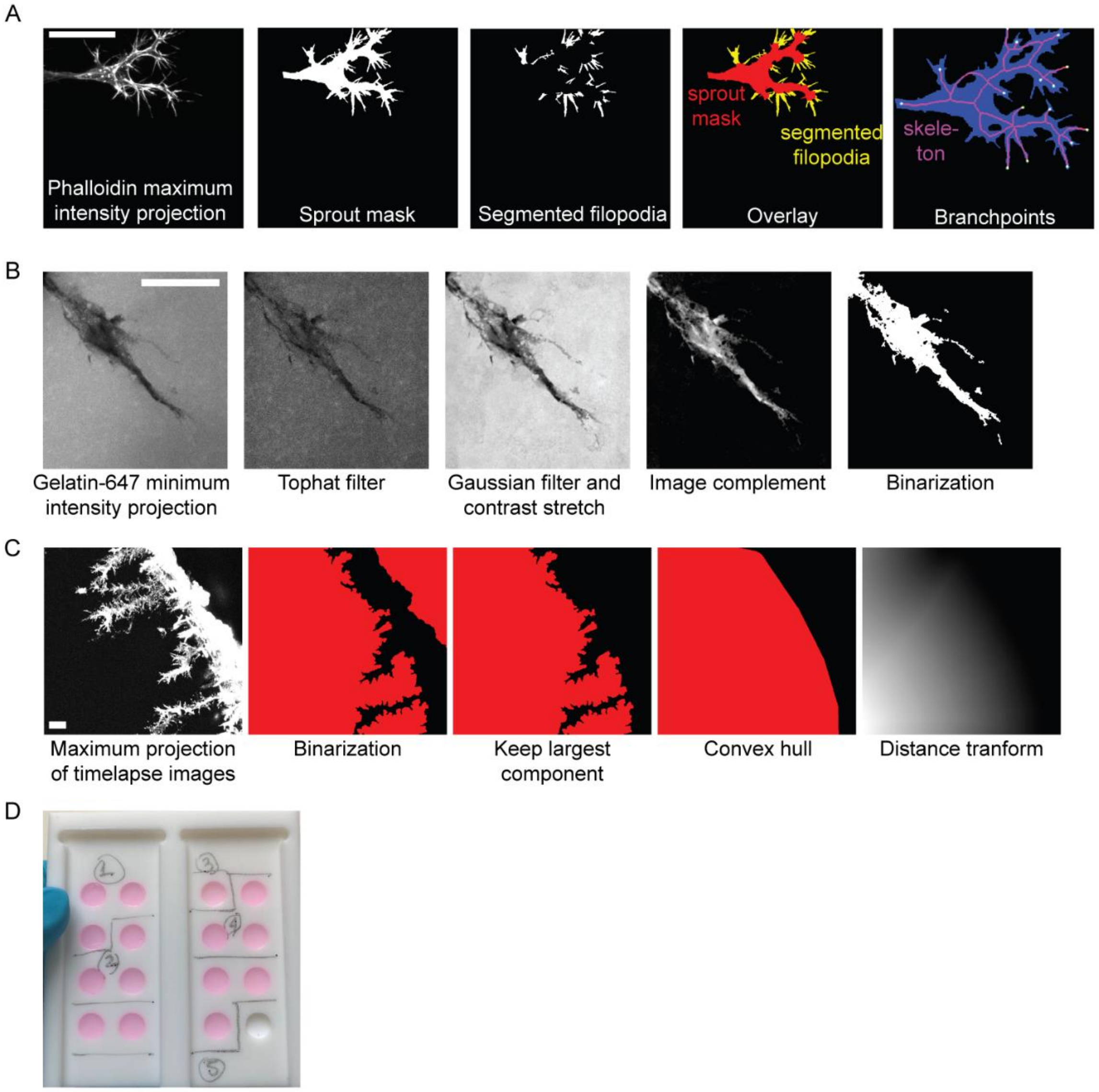

**Supplementary Fig. 1: Quantification methods. A.** Representative steps for quantification of filopodia. From left to right: maximum projection of an image stack of a phalloidin labelled sprout, binary mask obtain after thresholding, binary mask after subtraction of the mask processed with the image opening operation, overlay of both binary masks, and sprout mask (blue), skeleton (red), and detected branch points (green dots). **B.** Representative steps for quantification of matrix degradation. From left to right: minimum intensity projection of a 3D stack of PEG combined with the Gelatin-647 dye, image after removal of uneven background illumination with a top-hat filter, image after gaussian filtering and contrast stretching operations, image after computing its complementary, and the binary mask after intensity thresholding. **C.** Representative steps for automated quantification of dynamic invasion. From left to right: maximum intensity projection of the timelapse for one of the cell channels, binary mask of the complementary image, binary mask after retention of the largest component, convex hull of the binary image, and distance transform of the binary image. All scalebars are 50µm. **D.** Homebuilt mold used for sample preparation for unconfined compression testing.

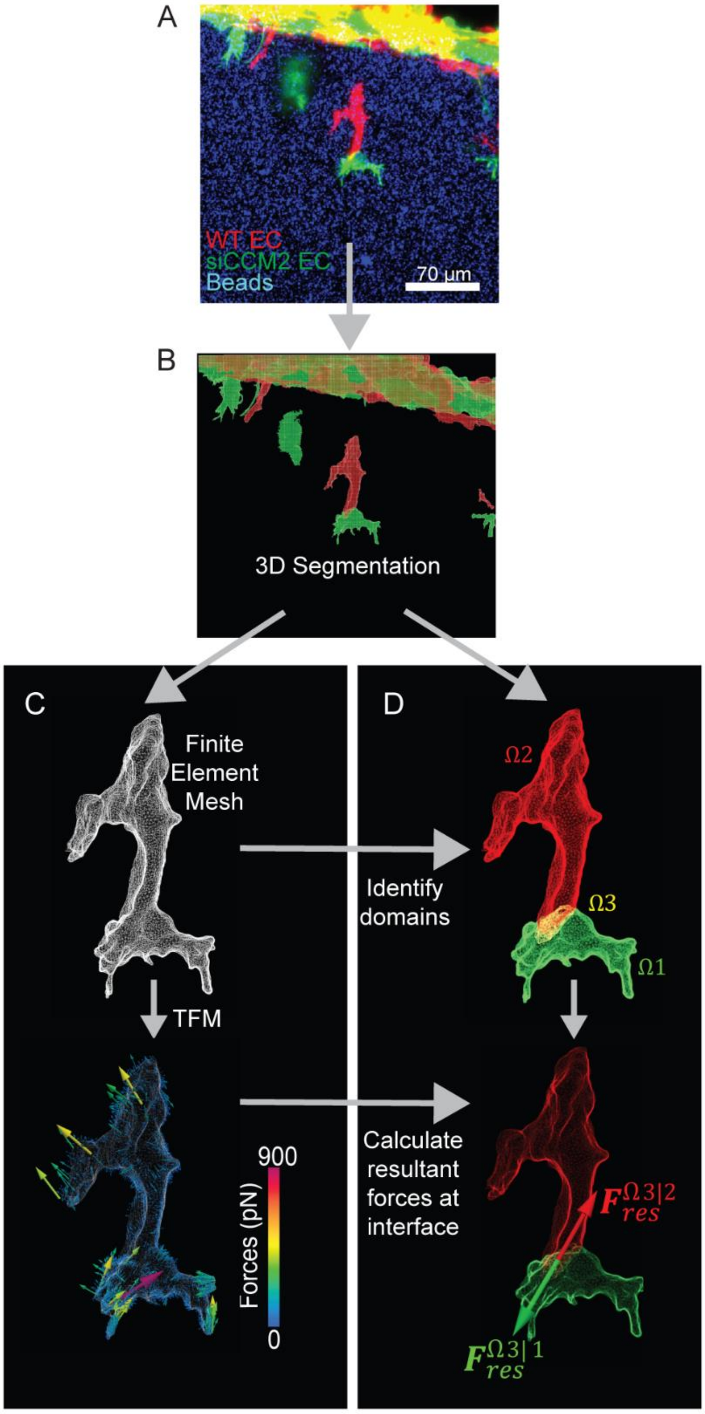

**Supplementary Fig. 2: Quantification of cell-cell forces.** a) Maximum intensity projections of the original confocal microscopy image of the cell pair. siCCM2 EC are labelled in green, WT EC in red, and the fluorescent beads embedded in the PEG gel, in blue. b) Overlay of the 3D segmentations (image voxel space) obtained through TFMLAB. c) Finite element mesh of the surface of the cell pair (top) and force distribution expressed in nN (bottom). d) Identification of the 3 domains (cell 1 in green, cell 2 in red, and cell-cell interface in yellow) in the finite element mesh by finding the nearest neighbors to the voxel coordinates of b) (top); representation of the $\boldsymbol{F}_{res}^{\Omega3|1}$ and $\boldsymbol{F}_{res}^{\Omega3|2}$ vector directions, confirming the force balance in the cell domain.

### Captions for Supplementary Videos

**Supplementary Video 1: Example of WT EC following tunnels created by leading mutant.** Timelapse imaging in a single z-plane of a trailing WT EC (in red) and leading siCCM2 mutant EC (in green) with tunnels (in black) seen in the gelatin embedded hydrogel behind the migrating siCCM2. Yellow arrow refers to timepoints in which the trailing WT EC tip modifies its direction to sense the tunnel.

**Supplementary Video 2A: Representative example of dynamic sprouting in control mosaics from Fig. 2c.** Maximally projected confocal timelapse ($\Delta t$=45min) images from sprouting of a 1:1 mix of WT ECs (in red) and siCT ECs (in green) into PEG hydrogel. Scale bar is 50 µm.

**Supplementary Video 2B: Representative example of dynamic sprouting in CCM2 mosaics from Fig. 2c.** Maximally projected confocal timelapse ($\Delta t$=45min) images from sprouting of a 1:1 mix of WT ECs (in red) and siCCM2 ECs (in green) into PEG hydrogel. Scale bar is 50 µm.

**Supplementary Video 2C: Representative example of dynamic sprouting in CCM2+R1 mosaics from Fig. 2i.** Maximally projected confocal timelapse ($\Delta t$=45min) images from sprouting of a 1:1 mix of WT ECs (in red) and siCCM2+SiR1 ECs (in green) into PEG hydrogel. Scale bar is 50 µm.

**Supplementary Video 2D: Representative example of dynamic sprouting in CCM2+R2 mosaics from Fig. 2i.** Maximally projected confocal timelapse ($\Delta t$=45min) images from sprouting of a 1:1 mix of WT ECs (in red) and siCCM2+siR2 ECs (in green) into PEG hydrogel. Scale bar is 50 µm.

**Supplementary Video 3: Representative example of reduced migration of stalk WT ECs upon losing contact with detached leading mutant EC.** Maximally projected confocal timelapse ($\Delta t$=50min) images from sprouting of 1:1 mix of WT ECs (in red) and siCCM2 ECs (in green). Images were zoomed, cropped and edited to remove background invasion in different planes for clarity of visualization. Unedited video is also available upon request. Scale bar is 50 µm.

**Supplementary Video 4: Representative example of the dynamic cell tracking algorithm.** (left) Timelapse ($\Delta t$=45min) of maximum intensity projections of confocal images of Lifeact-GFP labelled siCCM2 ECs invading a PEG gel. (right) Results from the individual cell tracking algorithm. Each color identifies one of the detected cells.

#### Captions for Supplementary Files

**File 1:** Top marker genes per cluster

**File 2:** Gene-sets used in Fig. 4d

**File 3:** Top 50 marker genes per cell type in Fig. 4e

**File 4:** Gene-sets used in Fig. 4h

### Supplementary Note 1: 3D Traction force microscopy data analysis

Given a FE mesh and measured displacements, this method searches for a displacement field that is as similar as possible to the measured one while satisfying the force equilibrium condition in the hydrogel domain.

Let $h$, $c$ and $b$ be the (interior) hydrogel, cell and (external) hydrogel boundary domains, respectively. Mathematically, this can be expressed as:

$$\min_{\boldsymbol{u}_c^{calc},\boldsymbol{u}_h^{calc}} \left(\frac{1}{2}\left\|\boldsymbol{u}_c^{calc}-\boldsymbol{u}_c^{meas}\right\|_2^2+\frac{1}{2}\left\|\boldsymbol{u}_b^{calc}-\boldsymbol{u}_b^{meas}\right\|_2^2\right) \quad \text{s.t.} \quad \boldsymbol{F}_h=0 \qquad \textit{Eq. 1}$$

where $\boldsymbol{F}$ represents the nodal reaction forces vector which accounts for external or internal prescribed forces or displacements. Including the equilibrium constraint as a Langrange multiplier, Eq. 1 can be rewritten as,

$$\min_{\boldsymbol{u}_c^{calc},\boldsymbol{u}_h^{calc}} \left(\frac{1}{2}\left\|\boldsymbol{u}_c^{calc}-\boldsymbol{u}_c^{meas}\right\|_2^2+\frac{1}{2}\left\|\boldsymbol{u}_b^{calc}-\boldsymbol{u}_b^{meas}\right\|_2^2+\Theta\cdot\eta\right) \qquad \textit{Eq. 2}$$

where Θ is the equilibrium constraint equation and η is the Lagrange multiplier. In a FEM framework, the discretized (and linearized) elasticity problem turns into the following algebraic system:

$$\begin{bmatrix}\boldsymbol{F}_c\\ \boldsymbol{F}_h\\ \boldsymbol{F}_b\end{bmatrix}=\begin{bmatrix}\boldsymbol{K}_{cc} & \boldsymbol{K}_{ch} & \boldsymbol{K}_{cb}\\ \boldsymbol{K}_{hc} & \boldsymbol{K}_{hh} & \boldsymbol{K}_{hb}\\ \boldsymbol{K}_{bc} & \boldsymbol{K}_{bh} & \boldsymbol{K}_{bb}\end{bmatrix}\begin{bmatrix}\boldsymbol{u}_c\\ \boldsymbol{u}_h\\ \boldsymbol{u}_b\end{bmatrix} \qquad \textit{Eq. 3}$$

Therefore, the optimization problem in Eq. 2 can be solved analytically and rewritten in matrix form as,

$$\begin{bmatrix}\boldsymbol{I} & \boldsymbol{0} & \boldsymbol{0} & \boldsymbol{K}_{ch}\\ \boldsymbol{0} & \boldsymbol{0} & \boldsymbol{0} & \boldsymbol{K}_{hh}\\ \boldsymbol{0} & \boldsymbol{0} & \boldsymbol{I} & \boldsymbol{K}_{bh}\\ \boldsymbol{K}_{hc} & \boldsymbol{K}_{hh} & \boldsymbol{K}_{hb} & \boldsymbol{0}\end{bmatrix}\cdot\begin{bmatrix}\boldsymbol{u}_c^{calc}\\ \boldsymbol{u}_h^{calc}\\ \boldsymbol{u}_b^{calc}\\ \eta\end{bmatrix}=\begin{bmatrix}\boldsymbol{u}_c^{meas}\\ \boldsymbol{0}\\ \boldsymbol{u}_b^{meas}\\ \boldsymbol{0}\end{bmatrix} \qquad \textit{Eq. 4}$$

where $\boldsymbol{K}_{hc}$, $\boldsymbol{K}_{ch}$ and $\boldsymbol{K}_{hh}$ are the parts of the stiffness matrix associated with the subdomains $c$, $h$ and $b$. TFMLAB makes use of the toolbox Suitesparse[8] to efficiently calculate $\boldsymbol{u}_c^{calc}$, $\boldsymbol{u}_h^{calc}$, $\boldsymbol{u}_b^{calc}$ and $\eta$. The values of $\eta$ are discarded since they do not hold mechanical meaning. Using a FEM framework, the strain field at each Gauss point of an element (e) is computed as,

$$\varepsilon^{(e)}=\boldsymbol{B}\cdot\boldsymbol{u}^{calc\,(e)} \qquad \textit{Eq. 5}$$

where $\boldsymbol{B}$ is the gradient matrix of the shape functions**.** The stress tensor is calculated at each Gauss point of an element $(e)$ based on the linear elastic, isotropic and homogeneous constitutive properties as,

$$\boldsymbol{\sigma}^{(e)}=2\mu\cdot\varepsilon^{(e)}+\lambda\cdot tr(\varepsilon^{(e)})\boldsymbol{I} \qquad \textit{Eq. 6}$$

where $\mu$ and $\lambda$ are the Lamé constants of the hydrogel. $\boldsymbol{\sigma}^{(e)}$ is then averaged from Gauss points to the nodes $i$ of the FE mesh. Tractions are finally computed at the cell boundary domain using the Cauchy relation,

$$\boldsymbol{T}_i=\boldsymbol{\sigma}_i\cdot\boldsymbol{n}_i \qquad \textit{Eq. 7}$$

where $\boldsymbol{T}_i$ is the traction vector at the nodes $i$ of the FE mesh and $\boldsymbol{n}_i$ is the outward normal to node $i$.

The hydrogel domain was modeled as a linear elastic material with E = 520 Pa (see hydrogel mechanical characterization section) and ν = 0.3. TFMLAB assumes a negligible stiffness for the cell internal domain (Young's modulus of $10^{-6}$ Pa).

To obtain the displacement and traction metrics displayed in Fig. 1g, h, we calculated the $90^{th}$ percentile of the displacement/traction magnitude at the surface nodes cell FE mesh.

### Supplementary Note 2: Cell-cell force quantification

We first selected a pair of cells from the chimera assay composed of a WT and a siCCM2 cell that had detached from the cell monolayer (Supplementary Fig. 2A). Then, since each cell is present in a different color channel, TFMLAB binarizes the images of both cell channels individually (Supplementary Fig. 2B) and then joins them to form a combined cell mask that can be used to generate a FE mesh (Supplementary Fig. 2C, top). While the forces that occur in the cell-cell contact region are not accessible in TFM, we developed a novel approach to quantify the resultant force at the cell-cell interface. First, we redefine the domain c, the cell domain, which includes now both WT and a siCCM2 cells. We then add additional constraints to Eq. 1Eq. 2 so that not only the equilibrium of forces is fulfilled in the hydrogel domain, $h$, but also that the sum of forces and moments (with respect to any prescribed point) in the cell domain is zero, as follows,

$$\begin{gathered}\min_{\boldsymbol{u}_c^{calc},\boldsymbol{u}_h^{calc}} \left(\frac{1}{2}\left\|\boldsymbol{u}_c^{calc}-\boldsymbol{u}_c^{meas}\right\|_2^2+\frac{1}{2}\left\|\boldsymbol{u}_b^{calc}-\boldsymbol{u}_b^{meas}\right\|_2^2\right)\\ s.t.\\ \boldsymbol{F}_h=0\\ \sum_i^{N_c}\boldsymbol{F}_c=0\\ \sum_i^{N_c}\boldsymbol{r}_c\times\boldsymbol{F}_c=0\end{gathered} \qquad \text{Eq. 8}$$

where $N_c$ is the number of nodes in the cell domain. The last equation shown in Eq. 8 denotes moment equilibrium of forces in $c$. Thus, $\boldsymbol{r}_c$ is the vector position of each node in c meaning that equilibrium of moments is taken from the origin of coordinates. The term $\boldsymbol{r}_c\times\boldsymbol{F}_c$ is further elaborated as follows (with $\boldsymbol{F}_c$ being forces exerted on the hydrogel by the cells):

$$\boldsymbol{r}_c\times\boldsymbol{F}_c=\boldsymbol{F}_c\cdot\mathbb{I} \qquad \text{Eq. 9}$$

with $\mathbb{I}$ being a matrix operator that includes the nodal positions of $c$, properly ordered to account for the vector product. Similarly, as in Eq. 2, we included the equilibrium constraints through Lagrange multipliers as follows:

$$\begin{aligned}\min_{\boldsymbol{u}_c^{calc},\boldsymbol{u}_h^{calc}} \Bigg(&\frac{1}{2}\left\|\boldsymbol{u}_c^{calc}-\boldsymbol{u}_c^{meas}\right\|_2^2+\frac{1}{2}\left\|\boldsymbol{u}_b^{calc}-\boldsymbol{u}_b^{meas}\right\|_2^2+\left(\boldsymbol{K}_{hc}\boldsymbol{u}_c^{calc}+\boldsymbol{K}_{hh}\boldsymbol{u}_h^{calc}+\boldsymbol{K}_{hb}\boldsymbol{u}_b^{calc}\right)\boldsymbol{\eta}\\ &+\left(\sum_i^{N_c}\boldsymbol{K}_{cc}\boldsymbol{u}_c^{calc}+\sum_i^{N_c}\boldsymbol{K}_{ch}\boldsymbol{u}_h^{calc}+\sum_i^{N_c}\boldsymbol{K}_{cb}\boldsymbol{u}_b^{calc}\right)\boldsymbol{\xi}\\ &+\left(\sum_i^{N_c}\mathbb{I}_{cc}\cdot\boldsymbol{K}_{cc}\boldsymbol{u}_c^{calc}+\sum_i^{N_c}\mathbb{I}_{cc}\cdot\boldsymbol{K}_{ch}\boldsymbol{u}_h^{calc}+\sum_i^{N_c}\mathbb{I}_{cc}\cdot\boldsymbol{K}_{cb}\boldsymbol{u}_b^{calc}\right)\boldsymbol{\varsigma}\Bigg)\end{aligned}$$

Eq. 10

Solving
Eq. 10 analytically and expressing it in matrix form, we obtain:

$$\begin{bmatrix} \boldsymbol{I} & \boldsymbol{0} & \boldsymbol{0} & \boldsymbol{K}_{ch} & \sum_{i}^{N_c} \boldsymbol{K}_{cc} \cdot \boldsymbol{1_c} & \sum_{i}^{N_c} \boldsymbol{K}_{cc} \cdot \mathbb{I}_c \\ \boldsymbol{0} & \boldsymbol{0} & \boldsymbol{0} & \boldsymbol{K}_{hh} & \sum_{i}^{N_c} \boldsymbol{K}_{ch} \cdot \boldsymbol{1_h} & \sum_{i}^{N_c} \boldsymbol{K}_{ch} \cdot \mathbb{I}_h \\ \boldsymbol{0} & \boldsymbol{0} & \boldsymbol{I} & \boldsymbol{K}_{bh} & \sum_{i}^{N_c} \boldsymbol{K}_{cb} \cdot \boldsymbol{1_b} & \sum_{i}^{N_c} \boldsymbol{K}_{cb} \cdot \mathbb{I}_b \\ \boldsymbol{K}_{hc} & \boldsymbol{K}_{hh} & \boldsymbol{K}_{hb} & \boldsymbol{0} & \boldsymbol{0} & \boldsymbol{0} \\ \sum_{i}^{N_c} \boldsymbol{1_c^T} \cdot \boldsymbol{K}_{cc} & \sum_{i}^{N_c} \boldsymbol{1_h^T} \cdot \boldsymbol{K}_{hc} & \sum_{i}^{N_c} \boldsymbol{1_b^T} \cdot \boldsymbol{K}_{bc} & \boldsymbol{0} & \boldsymbol{0} & \boldsymbol{0} \\ \sum_{i}^{N_c} \mathbb{I}_{\boldsymbol{c}}^{\boldsymbol{T}} \cdot \boldsymbol{K}_{cc} & \sum_{i}^{N_c} \mathbb{I}_{\boldsymbol{h}}^{\boldsymbol{T}} \cdot \boldsymbol{K}_{hc} & \sum_{i}^{N_c} \mathbb{I}_{\boldsymbol{b}}^{\boldsymbol{T}} \cdot \boldsymbol{K}_{bc} & \boldsymbol{0} & \boldsymbol{0} & \boldsymbol{0} \end{bmatrix} \cdot \begin{bmatrix} \boldsymbol{u}_c^{calc} \\ \boldsymbol{u}_h^{calc} \\ \boldsymbol{u}_b^{calc} \\ \boldsymbol{\eta} \\ \boldsymbol{\xi} \\ \boldsymbol{\varsigma} \end{bmatrix} = \begin{bmatrix} \boldsymbol{u}_c^{meas} \\ \boldsymbol{0} \\ \boldsymbol{u}_b^{meas} \\ \boldsymbol{0} \\ \boldsymbol{0} \\ \boldsymbol{0} \end{bmatrix}$$

Eq. 11

With **1** being the all ones vector. After solving $\boldsymbol{u}^{calc}$, nodal forces can be simply calculated with $\boldsymbol{F}^{calc} = \boldsymbol{K} \cdot \boldsymbol{u}^{calc}$ (see force distribution in Supp. Fig. 8c, bottom).

Then, since the FE mesh of the cell domain did not distinguish between the two cells of the cell pair, we used the voxel coordinates of the binary masks of each cell and, based on their nearest neighbor, we labelled each cell FE node accordingly. Moreover, the intersection between the two masks, i.e., the contact region between the two cells, was also labelled. Therefore, we split our FE mesh nodes in three domains: cell 1, $\Omega_1$, cell 2, $\Omega_2$, and the cell-cell interface region, $\Omega_3$ (see schematic of Fig. 3h and Supplementary Fig. 2D, top). We first analyzed which cell was mechanically more active by calculating the total force magnitude, $F_{tot}^{\Omega}$, of the cell-ECM force vectors, and the average traction magnitude, $T_{avg}^{\Omega}$, with the following expressions:

$$F_{tot}^{\Omega} = \sum_{i=1}^{N_\Omega} \|\boldsymbol{F}_i^{\Omega}\| \qquad \text{Eq. 12}$$

$$T_{avg}^{\Omega} = \frac{1}{A_{tot}^{\Omega}} \sum_{i=1}^{N_\Omega} \|\boldsymbol{F}_i^{\Omega}\| \qquad \text{Eq. 13}$$

Where $\|\blacksquare\|$ is the magnitude of a vector, $N_\Omega$ is the number of nodes in a given domain $\Omega$, and $A_{tot}^{\Omega}$ is its total surface area of domain $\Omega$. Results showed that cell-ECM tractions exerted by cell 1 (siCCM2 EC) were around 55% higher than those exerted by cell 2 (WT EC) (see values of $T_{avg}^{\Omega_1}$ and $T_{avg}^{\Omega_2}$ in Table 1).

Because each individual cell must also be in equilibrium of forces, the following expressions must hold:

$$\sum_{i=1}^{N_{\Omega_1}} \boldsymbol{F}_i^{\Omega_1} + \sum_{i=1}^{N_{\Omega_3}} \boldsymbol{F}_i^{\Omega_3} = 0; \ \boldsymbol{F}_{res}^{\Omega_{3|1}} = \sum_{i=1}^{N_{\Omega_3}} \boldsymbol{F}_i^{\Omega_3} = -\sum_{i=1}^{N_{\Omega_1}} \boldsymbol{F}_i^{\Omega_1} \qquad \text{Eq. 14}$$

$$\sum_{i=1}^{N_{\Omega_2}} \boldsymbol{F}_i^{\Omega_2} - \sum_{i=1}^{N_{\Omega_3}} \boldsymbol{F}_i^{\Omega_3} = 0; \ \boldsymbol{F}_{res}^{\Omega_{3|2}} = -\sum_{i=1}^{N_{\Omega_3}} \boldsymbol{F}_i^{\Omega_3} = -\sum_{i=1}^{N_{\Omega_2}} \boldsymbol{F}_i^{\Omega_2} \qquad \text{Eq. 15}$$

where $\boldsymbol{F}_i^{\Omega_1}, \boldsymbol{F}_i^{\Omega_2}, \boldsymbol{F}_i^{\Omega_3}, N_{\Omega_1}, N_{\Omega_2}, N_{\Omega_3}$ are the nodal forces and the number of nodes in the $\Omega_1$, $\Omega_2$, and $\Omega_3$ domains, respectively, with $\boldsymbol{F}_i^{\Omega_1}$ and $\boldsymbol{F}_i^{\Omega_2}$ as calculated by TFM (and therefore being forces exerted on the hydrogel by cell 1 and cell 2, respectively). $\boldsymbol{F}_{res}^{\Omega_{3|1}}$(Eq. 14) can be interpreted as the resultant force exerted by cell 1 on cell 2. Similarly, $\boldsymbol{F}_{res}^{\Omega 3|2}$ (Eq. 15) can be interpreted as the resultant force exerted by cell 2 on cell 1. Therefore, with the forces $\boldsymbol{F}_i^{\Omega_1}$ and $\boldsymbol{F}_i^{\Omega_2}$ at the cell-ECM boundary domain provided by TFM, it is possible to obtain the resultant force (sum of forces) at the cell-cell interface domain. In an error-free situation, the resultant forces $\boldsymbol{F}_{res}^{\Omega_{3|1}}$ and $\boldsymbol{F}_{res}^{\Omega_{3|2}}$ are equal with opposite sign. Indeed, our results showed that both resultant vectors point in opposite directions (Supplementary Fig. 2D, bottom) and have similar magnitudes (~14 nN; see Table 1). The fact that they are more than one order of magnitude larger than the computing error (which can be estimated from the difference between $\left\|\boldsymbol{F}_{res}^{\Omega_{3|1}}\right\|$ and $\left\|\boldsymbol{F}_{res}^{\Omega_{3|2}}\right\|$) is a first sign that cell-cell forces are non-negligible. Secondly, to understand whether this resultant force at the cell-cell interface has a magnitude that is comparable to that of cell-ECM forces, we calculated the average force magnitude per unit of area at the cell-cell interface domain. Because our approach only provides a resultant force at the cell-cell interface and not a distribution of forces, we divided the magnitude of the resultant force by the surface area of the cell-cell interface domain with the expressions:

$$T_{avg}^{\Omega_{3|1}} = \frac{1}{A_{\Omega 3}} \left\|\boldsymbol{F}_{res}^{\Omega_{3|1}}\right\| \qquad \text{Eq. 16}$$

$$T_{avg}^{\Omega_{3|2}} = \frac{1}{A_{\Omega 3}} \left\|\boldsymbol{F}_{res}^{\Omega_{3|2}}\right\| \qquad \text{Eq. 17}$$

Table 1 shows that, evidently, both $T_{avg}^{\Omega_{3|1}}$, $T_{avg}^{\Omega_{3|2}}$ are almost equal. Moreover, the results show that these tractions are in the same order of magnitude as the average cell-ECM tractions, $T_{avg}^{\Omega_1}$ and $T_{avg}^{\Omega_2}$ (ranging between 6 and 16 pN/μm$^2$). This confirmed that substantial forces take place at the cell-cell interface.

*Table 1: Metrics used to study force exertion in the cell-cell pair.*

| Metric | Value | Definition of the metric | Formula |
|---|---|---|---|
| $F_{tot}^{\Omega_1}$ | 170.37 nN | Total force magnitude exerted by the CCM EC on the ECM | Eq. 12 |
| $F_{tot}^{\Omega_2}$ | 165.61 nN | Total force magnitude exerted by the WT EC on the ECM | Eq. 12 |
| $T_{avg}^{\Omega_1}$ | 16.85 pN/μm$^2$ | Average traction magnitude exerted by the CCM EC on the ECM | Eq. 13 |
| $T_{avg}^{\Omega_2}$ | 9.44 pN/μm$^2$ | Average traction magnitude exerted by the WT EC on the ECM | Eq. 13 |
| $\left\|\boldsymbol{F}_{res}^{\Omega_{3\mid1}}\right\|$ | 14.70 nN | Magnitude of the resultant force exerted by the CCM EC on the WT EC | Eq. 14 |
| $\left\|\boldsymbol{F}_{res}^{\Omega_{3\mid2}}\right\|$ | 13.90 nN | Magnitude of the resultant force exerted by the WT EC on the CCM EC | Eq. 15 |
| $T_{avg}^{\Omega_{3\mid1}}$ | 7.22 pN/μm$^2$ | Average traction magnitude exerted by the CCM EC on the WT EC | Eq. 16 |
| $T_{avg}^{\Omega_{3\mid2}}$ | 6.79 pN/μm$^2$ | Average traction magnitude exerted by the WT EC on the CCM EC | Eq. 17 |

Given the fact that the resultant cell-cell forces $\boldsymbol{F}_{res}^{\Omega_{3|1}}$ and $\boldsymbol{F}_{res}^{\Omega_{3|2}}$ point away from the respective cell on which they act, we conclude that cells apply pulling (instead of pushing) forces on each other. Moreover, we can compare the direction (and sign) of $\boldsymbol{F}_{res}^{\Omega_{3|1}}$ (i.e., the cell-cell force on the WT follower EC) to the direction of invasion of the cell pair and assess its importance for the motion of the WT follower EC. To this end, we obtained the direction of invasion of the cell pair by defining a vector, $\boldsymbol{m}$, whose direction pointed from the centroid of the cell monolayer to the centroid of the leading cell (see blue arrow in Fig. 3h). As $\boldsymbol{F}_{res}^{\Omega_{3|1}}$ and $\boldsymbol{m}$ point in the same direction (i.e. the dot product $\boldsymbol{F}_{res}^{\Omega_{3|1}} \cdot \boldsymbol{m}$ is positive) the cell-cell resultant force is likely to be driving the motion of the WT follower EC (and not the resultant cell-ECM force on the follower cell, which points in the opposite direction).

At the level of the cell pair (for which cell-cell forces are internal and therefore do not contribute to its motion), cell-ECM pulling forces exerted by the leading siCCM2 are stirring cell pair migration. Given the higher mechanical activity of the siCCM2 leader (see also average cell-ECM tractions in Table 1) we assume that cell-cell pulling forces are mainly originating from the siCCM2 leader's (internal) contractile machinery.
